# Protein Crowders Remodel RNA Electrostatics, Hydration, and Dynamics: A Challenge to Steric Crowding Models

**DOI:** 10.1101/2025.09.25.678182

**Authors:** Anja Henning-Knechtel, Marko Brnović, Weiwei He, Serdal Kirmizialtin

## Abstract

The intracellular environment is densely populated with macromolecules, creating crowded conditions. Whether in vitro environments or synthetic crowders like polyethylene glycol (PEG) accurately capture the complexity of RNA interactions in vivo remains unclear. Using extensive all-atom molecular dynamics simulations, we investigated the HIV-1 TAR RNA hairpin in dilute, PEG-crowded, and realistic protein-crowded solutions. We found that PEG primarily exerts excluded-volume effects, maintaining RNA hydration and Na^+^ ion condensation similar to dilute conditions. In contrast, protein crowders significantly altered RNA electrostatics, reducing Na^+^ condensation by nearly 60%, displacing surface hydration water, and forming chemically specific contacts dominated by positively charged residues, notably arginine and lysine. These interactions led to local RNA expansion and reshaped its conformational landscape without disrupting the global fold. Moreover, protein crowding dramatically slowed RNA translational and rotational dynamics, as well as local water and ion mobility, effects minimally observed with PEG. Our findings emphasize that crowder identity critically determines RNA behavior and challenge the use of PEG as a universal model for intracellular conditions, providing mechanistic predictions for RNA studies in biologically relevant environments.

## Introduction

RNA functions within the highly crowded and complex cellular environment, significantly impacting its folding, interactions, and dynamics; however, the precise effects of crowding on RNA electrostatics and hydration remain incompletely understood. The intracellular environment is a crowded, compositionally complex, and densely populated with macromolecules, including proteins, RNAs, DNA, polysaccharides, and metabolites, reaching concentrations of up to 400mg/mL in the bacterial cytosol and 100–300mg/mL in eukaryotic cells^1–4^. This molecular crowding exerts profound effects on the structure, dynamics, and function of biomolecules, not only through excluded-volume effects but also via direct physical and chemical interactions, altering thermodynamic and kinetic properties ^5,6^. While crowding effects on protein folding and compaction have been extensively characterized ^5,7–9^, the impact on nucleic acids, particularly RNA, is limited to folding and binding stability^10–12^, many of the properties of the solvation remain to be understood ^13^.

RNA presents unique challenges and opportunities for crowding studies due to its polyanionic backbone, high hydration demand, and ion-dependent tertiary structure formation. Unlike DNA, which readily condenses in response to crowding and multivalent ions^14–17^, RNA adopts compact yet conformationally dynamic structures even under moderate crowding^10,18^. Studies have shown that crowding enhances RNA folding by favoring compact conformations via entropic stabilization ^10,19^, alters folding landscapes ^20^, and reduces dependence on divalent cations like Mg^2^+ for tertiary stabilization ^20–22^. Macromolecular crowding stabilizes the RNA by favoring its compact tertiary structure, tightening the ligand-binding pocket, and shifting the folding equilibrium toward the functional states. ^23,24^ However, RNA is also highly sensitive to electrostatic and hydration-mediated forces, and the precise mechanisms by which crowding modulates these forces remain unresolved ^25,26^. Classical counterion condensation theory^27^ explains ion-RNA interactions in dilute solution, but it fails to prediction behavior under confinement or altered dielectric conditions. Statistical mechanical and tightly bound ion models ^28^ have begun to incorporate these factors, but still require validation from explicit-solvent simulations.

Existing theoretical, computational, and experimental studies have outlined several competing effects of crowding on RNA electrostatics. First, excluded volume enhances folding stability by entropically favoring compact states ^10,19,20^. Second, crowding can alter the local dielectric environment, particularly through dehydration, potentially increasing electrostatic interactions ^29–31^. This arises in part from the dielectric decrement effect, where water structuring near macromolecular surfaces lowers local permittivity, amplifying electrostatic fields. Third, crowding agents, especially biological crowders such as proteins, can interact directly with the RNA surface, potentially displacing counterions and perturbing the ion atmosphere ^12,32–34^. Statistical mechanical models predict that such interactions can reduce counterion condensation and modulate the folding free energy landscape ^19,28^, yet these predictions have lacked direct atomistic validation. Coarse-grained models have demonstrated that RNA folding is modulated by crowding, highlighting the importance of explicitly accounting for ionic and hydration effects ^19^. While coarse-grained models have qualitatively suggested that RNA folding can be modulated under crowding, the explicit atomistic detail provided here is critical for accurately capturing ionic, electrostatic, and hydration effects that coarse-grained approaches typically oversimplify.

Moreover, synthetic crowders such as polyethylene glycol (PEG) are frequently used to mimic cellular crowding in vitro. PEG is attractive due to its inert nature and tunable molecular weight, but lacks molecular anisotropy, electrostatic surfaces, and the capacity to form structured hydration shells, all of which are critical for RNA-crowder interactions in vivo. The assumption that PEG adequately models cellular crowding has rarely been tested at the level of electrostatic and hydration perturbations. However, PEG lacks chemical complexity and heterogeneity, raising concerns about its ability to replicate the physical and electrostatic features of the intracellular environment ^11,33,35^. Studies on proteins or DNA generally regard PEG as a predominantly steric crowding agent, whereas biological crowders involve soft interactions and heterogeneous charge distributions that affect hydration and electrostatics in more intricate ways ^35–39^. In RNA, this distinction is particularly critical given the central role of ion–RNA interactions in tertiary structure stability and functional dynamics ^40,41^.

Despite the recognized importance of these issues, direct atomistic insight into how crowding, particularly from protein components, affects the electrostatic landscape, hydration shell, and ion association of RNA remains limited. Previous MD studies have resolved hydration dynamics and counterion behavior in dilute RNA systems or DNA, but few have tackled the full complexity of crowded RNA systems using explicit-solvent, atomistic simulations with realistic crowder compositions.

In this work, we address this gap by performing long-timescale, all-atom molecular dynamics simulations of the HIV-1 TAR RNA hairpin in three physiologically relevant environments: (i) a dilute aqueous NaCl solution, (ii) a PEG-crowded solution representing synthetic crowding, and (iii) a protein-crowded solution constructed from a mixture of cytoplasmic and cytoskeletal proteins at realistic intracellular concentrations. These simulations allow us to examine, at atomistic resolution, how different crowders influence RNA salvation, ion distribution, hydration shell dynamics, and electrostatic potential.

We find that while PEG exerts predominantly steric effects, preserving a Na^+^ coordination environment similar to dilute conditions, protein crowders profoundly perturb RNA electrostatics, displacing counterions and restructuring hydration shells through non-specific interactions, particularly involving positively charged residues. The resulting effect is that RNA is compressed in PEG while it is expanded in protein crowder. Counter-ion condensation theory failed to describe the electrostatic screening, where we observe more than two-fold decrease in the condensed counter ions on the RNA surface in protein crowders, while PEG crowders remain similar to the dilute RNA regime. These effects are not captured by simple volume exclusion models and suggest that protein crowding reshapes the RNA’s energetic landscape via a combination of electrostatic modulation, hydration layer displacement, and direct surface interactions.

By comparing synthetic and biological crowding agents, our study provides atomically detailed insight into the limits of PEG-based crowding mimetics and underscores the need to consider crowder identity and interaction specificity when modeling RNA folding and function in vitro. Our results serve as a benchmark for future simulation and experimental efforts aimed at capturing RNA behavior under native-like cellular conditions.

## Methods

All-atom MD simulations were carried out with GROMACS 2028.8 to investigate the structure and dynamics of RNA in different environments. The AMBER force field was used for the protein, supplemented with CUFIX corrections for protein-RNA interactions ^42^, while χOL3 force field^43^ was applied to RNA. Na^+^ ions were represented by Joung-Cheatham parameters44, and the TIP4P water model^45^ was employed. PEG 6000 was chosen to match commonly used crowding experiments. Parameters for PEG 6000 were generated using the Scienomics MAPS software suite 4.4.0. The investigations considered one simulation of the RNA in an aqueous NaCl solution and two simulations of the RNA in the presence of PEG or protein crowders. Different analyses were performed to identify the structural and dynamic differences of RNA in the three different environments.

### Preparation of the simulation box

To explore the influence of various environments on RNA behavior, we selected the transactivation response element (TAR) RNA from the human immunodeficiency virus 1 (HIV-1) as our model system, referred to as HIV-1 TAR RNA. This RNA molecule is widely employed in biophysical studies due to its relevance to HIV and its complex structure, encompassing stem regions, bulges, and an apical loop. HIV-1 TAR RNA is also recognized for its interaction with the TAT protein, facilitated by forming an arginine sandwich motif within the TAR bulge region ^46–50^. Further details on the preparation of the coordination file are provided in the Methods section of reference ^51^.

This refined RNA structure was positioned at the center of a triclinic simulation box with a volume of 4372.95 *nm*^3^ as a starting point for the subsequent setups. In the first simulation setup, modeling a standard non-crowded *in-vitro* experiment, explicit water molecules and NaCl were added to neutralize the RNA charges and to replicate common *in-vivo* and *in-vitro* experimental concentrations of 150 mM. Despite the predominance of *K*^+^ ions at a concentration of approx. 140-150 mM as the primary intracellular monovalent ion species^52^, we opted for *Na*^+^ in our experiments due to its prevalence in *in-vitro* experiments ^12^. We intentionally excluded *Mg*^2+^ to isolate the electrostatic and hydration effects of monovalent ions under crowding conditions. Because, unlike *Na*^+^, which interacts diffusely with the phosphate backbone, *Mg*^2+^ binds tightly to specific RNA motifs and could obscure the generalizable features of crowder-mediated ion exclusion ^40,41,53^. Additionally, including *Mg*^2+^ would introduce site-specific binding complexities, potentially obscuring more generalizable electrostatic crowding effects that we aimed to elucidate with monovalent ions. The Mg-RNA interactions in the crowding environment will be published later. The same salt concentration was utilized in all simulation boxes to maintain consistency.

Two PEG-crowding simulation boxes were prepared by randomly placing 72 PEG 6000 molecules around the centered RNA before adding water molecules and ions. The resulting crowder concentration of 165.22 g/l, falling within the reported macromolecular concentration range in a eukaryotic cell’s cytoplasm ^2,35,54^. PEG 6000 parameters were benchmarked against experimental density and radius of gyration in aqueous solution, consistent with prior PEG MD studies ^39^. Protein crowders were modeled using standard Amber parameters with full sidechain flexibility. Mimicking the cytoplasm protein composition, we created two protein-crowding simulations by inserting copies of four cytosolic and microtubule proteins that are highly expressed in various human cell lines ^55^. Specifically, eight ribosomal proteins S18 (S18, AlphaFoldDB: P62269), five translation-controlled tumor proteins (TP, AlphaFoldDB: P13693), three ribosomal proteins L23 (L23, AlphaFoldDB: P62829), and three tubulin alpha lb (Tα, AlphaFoldDB: P68363) were all positioned at least 1 nm away from the surface of RNA. These proteins span a range of sizes and net charges and are abundant in cytosolic and cytoskeletal fractions, ensuring a physiologically relevant representation while maintaining computational tractability. The resulting protein concentration of the system was found to be 164.97 g/1, which was similar to the PEG-crowding concentration.

### Molecular dynamics simulations

After preparation of the simulation systems, each box underwent energy minimization to refine the positions of solvent and solute atoms. This process was followed by a 1 ns-long solvent (NVT) and volume (NPT) simulations at 310 K, during which only ions and water molecules were allowed to move freely while the solute atoms were restrained to their initial positions using harmonic restraints with a force constant of 1000 *kJmol* ^− 1^*nm*^− 2^.

Subsequent simulation steps were carried out to equilibrate the individual molecular components. The initiation of each step was based on the dimensions and positions of the atoms in the last snapshot of the previous step. Initially, a 200-ns NPT simulation was performed to facilitate ion condensation onto the RNA and crowder molecules. The atoms of the solute components were harmonically restrained using a force constant of 1000 *kJmol*^− 1^*nm*^− 2^, while ions and water were allowed to move freely. Following this, an additional 200-ns long NPT simulation was conducted by removing the restraints on the proteins or PEG molecules, leaving only the RNA immobilized. The ion equilibration step was repeated in the case of the non-crowding RNA setup.

After these equilibration steps, all restraints were removed, and a production run in the form of an NPT simulation at 310 K was initiated. The crowder-free simulation ran for 1 *µ*s, while simulations for the PEG and protein environments lasted for 2 *µ*s. For each crowded condition, two independent simulations were run. Data was recorded every 5 ps for subsequent analysis.

### General Setting for Molecular Dynamics Simulations

All minimizations were conducted using the steepest descent method. Periodic boundary conditions were applied in three directions for all NVT and NPT simulations. Particle Mesh Ewald (PME) summation was employed to compute long-range electrostatic interactions, utilizing a 0.16-nm grid for the Fourier space summation, cubic interpolation for charge density on the grid, and a 1.0-nm real space distance cutoff for both electrostatics and van der Waals energies. A dispersion correction was applied for the van der Waals cutoff. Covalent bonds within water, RNA, protein, and PEG molecules were constrained to their equilibrium geometries using SETTLE^56^ and LINCS^57^ algorithms. The equations of motion were integrated using the Leap-Frog scheme with a time step of 1 fs. A velocity-rescaling thermostat^58^ was employed to maintain the temperature at 310 K. For NPT simulations, the pressure was set to 1 bar using a Parrinello-Rahman barostat ^59^.

### Data Analysis

#### Crowder Density Distribution

To characterize the crowding environment around the HIV-1 TAR hairpin, we computed spatial density distributions of crowder densities (proteins or PEG molecules) across three planar slices perpendicular to the RNA’s long axis (Z-axis). The RNA was first centered and aligned such that its helical axis coincided with the Z-direction, and both translational and rotational motions were removed throughout the trajectory.

For each slice, a slab of thickness Δ*z* = 4 Å along the Z-axis was selected, and the positions of crowder particles within the slab were projected onto the corresponding X-Y plane. This plane was discretized into a regular grid with cell dimensions Δ *x* × Δ*y*, and the local density *ρ*_*ij*_ in each grid cell (*i, j)* was computed as:

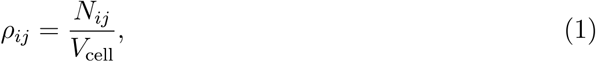

where *N*_*ij*_ is the number of crowder atoms within cell (*i, j)*, and *V*_cell_ = Δ *x* × Δ*y* × Δ *z* is the cell volume. For all slices, we used Δ*z* = 4 Å.

To facilitate comparison across different slices and crowding conditions, all computed densities are reported as raw values on each spatial grid. This analysis was repeated for three distinct regions along the RNA, providing spatially resolved insight into local crowding heterogeneity.

### Macromolecular size and scaling

To understand the crowder’s size and properties of the crowding environment, we used Flory’s scaling law to identify how crowders and solvents interact with each other. The Flory exponent *v* quantifies this behavior and is diagnostic of solvent quality: *v* ≈ 0.5 indicates that the crowder is in a theta solvent (ideal chain), *v* > 0.5 a good solvent, and *v* < 0.5 a poor solvent. This framework allows us to distinguish whether PEG and protein crowders form homogeneous or spatially heterogeneous environments for the RNA molecule at the given crowder concentration.

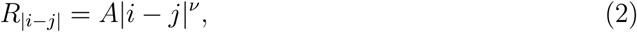

where |*i* − *j*| denotes the sequence separation between residues within the crowder, *v* is the scaling exponent, and *A* is a scaling prefactor. The scaling coefficient characterizes whether the solvent is good or poor for the macromolecule.

### Spectral Clustering of RNA Conformational Ensemble

To classify RNA conformations and compare structural ensembles across the different environments, we employ spectral clustering following method ^60^. For this analysis, RNA structures from all simulations were merged and aligned, and the pairwise root-mean-square deviation (RMSD) was calculated using the MDTraj library^61^:

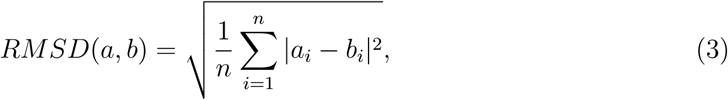

where *n* is the number of atoms in each structure, and *a*_*i*_, *b*_*i*_ denote the positions of atom *i* in structures *a* and *b*, respectively. This results in an *N* × *N* matrix of pairwise RMSD values across *N* RNA conformations.

Structures with an RMSD < 13.0 Å are considered similar. The RMSD matrix is converted into a binary adjacency matrix, where entries indicate whether two structures are within the similarity threshold. This adjacency matrix is used to construct a graph representation of RNA conformational space, which is then processed using spectral clustering. K-means analysis is employed for spectral embedding using scikit-learn ^62^.

### Local density analysis

The distribution of ions and crowder molecules around the RNA was investigated by radial distribution function (RDF), *g*_*X*−*Y*_(r). Here, *r* is the distance between the group *X* (ions or atoms of the crowder molecules) and the atoms of the group *Y* (RNA).

The surface of the RNA is divided into the major groove, the phosphate backbone, and the minor groove. Major groove is based on the nucleobase atoms N7, O6, N6, N4, and O4. The phosphate backbone is represented by OP1, OP2, and P, and the minor groove is represented by N3, N2, and O2 atoms on the RNA.

The cumulative number of bound ions on the RNA surface (*S*) up to distance *R, N*_*X*−*S*_(*R*), is computed from the RDF as

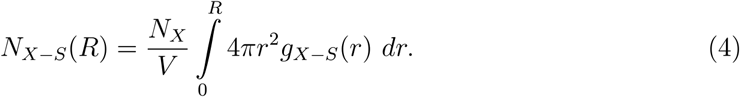

Here, *N*_*X*_ represents the total number of X in the simulation box, and *V* denotes the simulation box volume.

### Calculation of Electrostatic Properties from MD Simulations

To characterize the electrostatic environment around the RNA molecule from our molecular dynamics (MD) simulations, we computed the electric flux and electrostatic potential explicitly as functions of the distance *d*, measured from the RNA molecular surface outward.

We defined the RNA surface explicitly from the atomic positions obtained from our MD simulations, considering each atom’s van der Waals radius. At each distance *d* from the RNA surface, we considered a spherical region defined by extending this surface radially outward by distance *d*.

The total charge *Q*_enc_(*d*) enclosed within this spherical volume, comprising RNA, ions, and water molecules, was computed by summing the charges *q*_*i*_ of all atoms whose positions satisfy the condition:

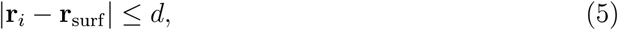

where **r**_*i*_ are atomic positions, and **r**_surf_ is the closest point on the RNA surface to the atom *i*.

Using Gauss’s law, the electric flux Φ_*E*_(*d*) through this surface at a distance *d* from the RNA was explicitly calculated as:

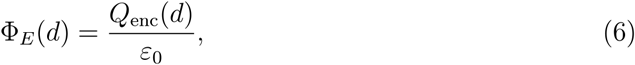

where *ε*_0_ is the vacuum permittivity (8.854 × 10^−12^ C^2^N^−1^m^−2^).

To complement the flux analysis, the electrostatic potential *V* (*d*) was computed explicitly by summing Coulombic contributions from each atom (RNA, ions, water molecules) enclosed within the spherical volume of radius *d* from the RNA surface. This potential was given by:

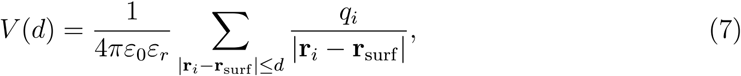

where *ε*_*r*_ is the relative permittivity (dielectric constant) reflecting the local dielectric environment provided by surrounding ions and water molecules. The values were computed for each time frame and averaged over the simulation time to obtain thermodynamically accessible observables.

### Contact Map analysis

Contact map analysis was used to compare conformations sampled at the different environments. Namely, the time average contact formation probability *C*(*i, j*) between residue pairs (*i*,*j*) with |*i* − *j*| > 2 was calculated as:

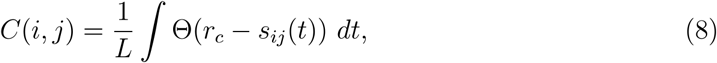

where Θ (*x*) is the Heaviside function, *L* is the simulation length, and *s*_*ij*_(*t*) is the shortest distance between the pairs in time. For our study, *r*_*c*_ was set to 6 Å to denote a contact. We compute the contacts between RNA -RNA and RNA and RNA-protein using the above criteria.

### Radius of gyration analysis

*R*_*g*_ was computed based on the eigenvalues of the tensor of inertia *λ*_*i*_ (*i*=l,2,3) according to:

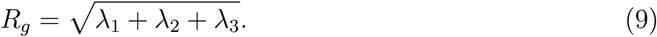

using the GROMACS plugin polystat.

### Residue and backbone flexibility analysis

Residue-wise fluctuation in atom positions was quantified by calculating the root-mean-square fluctuation (RMSF) by implementing the rmsf plugin of GROMACS. For this, the trajectory was fitted to the reference structure using a least-squares superposition of atomic coordinates, and the deviation of atom position over the trajectory was calculated as

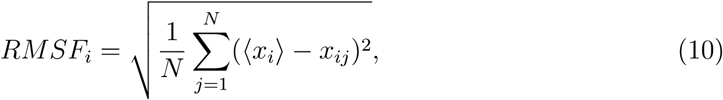

Where ⟨*x*_*i*_⟩ is the average position of the atom *i* over the trajectory length *N*, and *x*_*ij*_ is the position of the atom *i* in the frame *j* of the trajectory.

Furthermore, the backbone flexibility was assessed by determining the angle fluctuation between the C2-carbon of a nucleotide *i*, the phosphorus atom, and the C2-carbon atom of the adjacent nucleotide *i*+1 (C2^*i*^_P^*i*^_C2^*i*+l^ ). This angle reflects the mobility of the 2’-hydroxyl group, whose accessibility serves as a measure of backbone mobility in SHAPE reactivity experiments^63^. The fluctuation characterizes the effective stability, *ΔG*_*SIM*_, as:

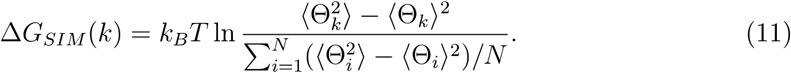

Here, the stability of the nucleotide *k* is computed from the fluctuation of this angle, where ⟨… ⟩ represents the ensemble average computed from the time series of the simulation trajectory. The value is normalized with the average fluctuation of *N* nucleotides of the RNA molecule.

The computed reactivity *a*_*SIM*_ can then be obtained as follows:

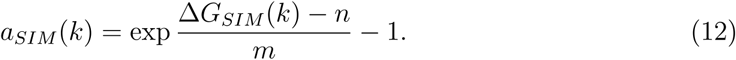

Fitting parameters *m* and *n* were set to 2.6 kcal/mol and 0.8 kcal/mol, respectively, following values reported for the E.coli 16S ribosomal RNA SHAPE data^64^.

### Hydrogen bond analysis

Hydrogen bond formation in a crowded and non-crowded environment was analyzed over the trajectory using the GROMACS-plugin hbond with a 3.5Å-cutoff.

### Solvent dynamics analysis

Investigation of the crowder influence on the dynamics of the RNA molecule and local environment included the lateral diffusion constant of the RNA and solvent components, rotational relaxation of the RNA movement, the structural correlation of the RNA, as well as the correlation of the number of cations and water molecules around the RNA.

The translational diffusion coefficient is defined as the derivative of the mean-square displacement with respect to time:

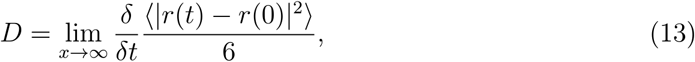

Where *r(t*) is the position of the geometric center of the molecule at time *t*. In this study, the diffusion constant for RNA, *Na*^+^ ions, and the oxygen atom of water was calculated from the slope of a straight-line fit in a 10-40 ns time window.

The RNA’s rotational orientation dynamics were quantified based on the orientational time-correlated function (*C*_*p*_(*t)*). This function relies on the selection of three specific atoms, namely the 05’ atom of residue lG, and the phosphate atom of 29G and 37C, to construct two vectors. The cross product of these vectors, denoted as **p**, captures the rotational dynamics of the RNA throughout the trajectory. The rotational autocorrelation function, *C*_*P*_, is then calculated based on the angular difference between the orientation of the cross product vector **p** at time. Namely, *C*_*P*_(*τ*) is defined as follows:

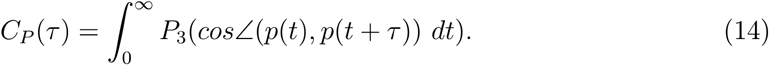

where *P*_3_(*x*) is a third order Legendre polynomial, *τ* is the elapsed time between the time points *t*. Higher order in Legendre polynomial allows capturing orientational dynamics of non-symmetric molecules. The integral over time averages the orientation correlation over the whole trajectory. This correlation function can be measured experimentally using techniques like NMR or dielectric relaxations.

Similarly, the correlation of the RNA structure, the ion distribution around the RNA, and the RNA hydration shell over time in crowded and non-crowded environments was computed based on an auto-correlation function:

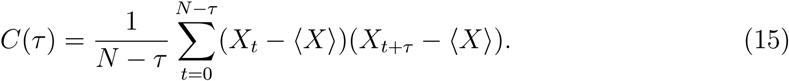

Here, *N* represents the total number of data points of the property *X*. For structural correlation, *X* corresponds to *R*_*g*_. For the correlation of the local RNA environment, *X* represents the number of cations or water molecules for specific distances from the RNA surface.

## Results

To examine how macromolecular crowding affects RNA structure and its electrostatic environment, we conducted all-atom molecular dynamics (MD) simulations. We performed multiple microsecond-long simulations of the HIV-1 TAR hairpin RNA under three distinct conditions: (i) a dilute, crowder-free *in vitro* solution (control, N), (ii) a PEG-crowded environment (G), and (iii) a protein-crowded environment (P) (see Figures 1 and S1). These simulations enabled a comparative analysis of how different crowding conditions influence the RNA’s structural dynamics and solvation. HIV-1 TAR RNA was chosen as a model system due to its diverse structural features, including 4-bp, 5-bp, and 12-bp duplex segments, a 6-nt apical loop, and 1-nt and 3-nt bulges. Further details are provided in the Methods section under **Preparation of the simulation box**.

**Figure 1:**
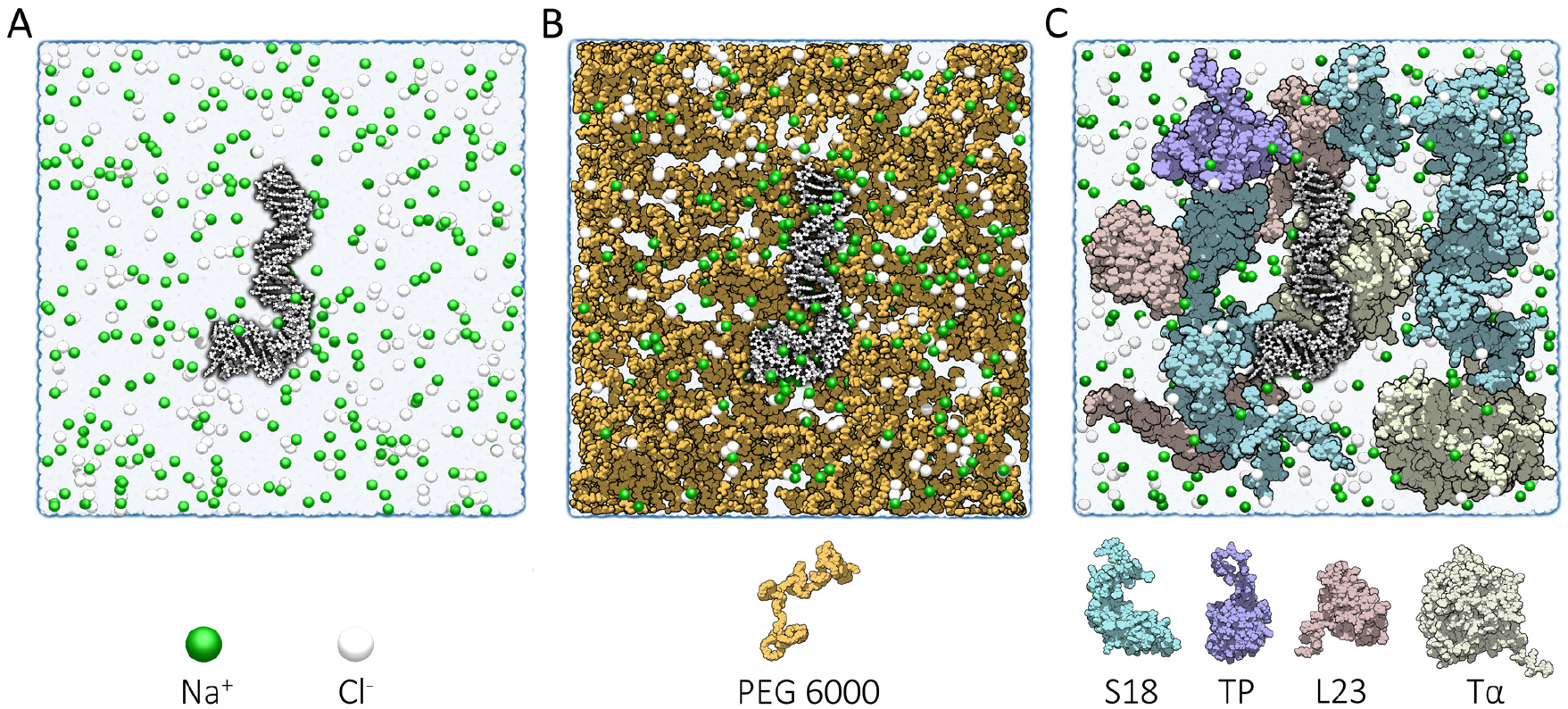
Cross-sectional view of the simulation box containing the RNA hairpin under different crowding conditions: (A) no crowders, (B) PEG crowders, and (C) protein crowders. Protein crowders were selected based on their expression levels across different cell lines. The specific protein crowders and ions are identified in the legend below the simulation setup. The cross-sections are taken after energy minimization along the x-axis (A, C) or y-axis (B) to align with the RNA orientation.

### Crowder Identity Dramatically Alters RNA Ionic Environment

We investigated how the surrounding environment influences the distribution of cations around the RNA molecule, focusing on sodium ions (Na^+^) commonly used in *in vitro* experiments. Figure 2A presents a snapshot from each setup with the cations within 6 A of the RNA shown in beads. The binding modes show a similarity between the *in vitro* (non-crowded) and PEG conditions. However, protein crowders lead to a depletion of the counterion atmosphere around the RNA.

**Figure 2:**
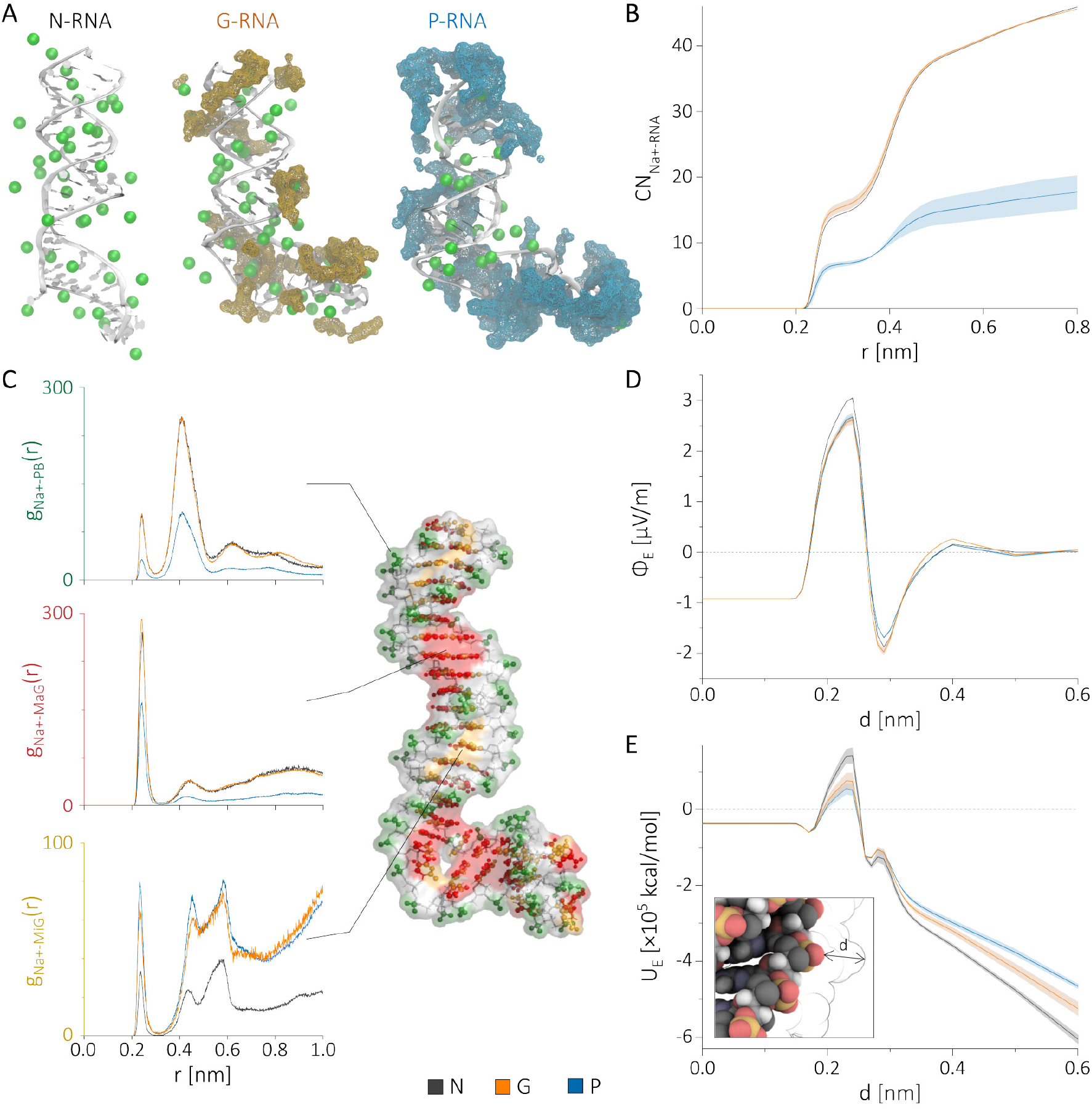
Environmental influences on RNA ionic distribution. (A) Snapshots illustrating the significant depletion of Na^+^ ions (green) around RNA (white) under protein crowding compared to PEG and non-crowded conditions. (B) Na^+^ coordination number as a function of distance from the RNA surface. (C) RDF profiles highlighting differential ionic interactions with RNA grooves. (D, E) Electric flux (Φ_*E*_) and potential energy (*U*_*E*_), illustrating crowder-dependent electrostatic compensation. Protein crowders show a substantial decrease in ion condensation compared to PEG and dilute conditions, altering RNA electrostatic stability.

To quantify these differences, we calculated the coordination number of cations relative to the RNA surface (Figure 2B). Without crowding, 41 Na^+^ ions are coordinated within 6 Å, resulting in nearly 80% charge neutralization. A similar level of charge compensation is observed in the PEG-crowded environment. Protein crowders reduce cation coordination to approximately 16 ± 2 Na^+^ ions, likely due to steric occlusion and electrostatic competition from positively charged residues at the RNA surface, leading to only 31% charge neutralization. Conversely, the coordination of anions (Cl^−^) (Figure S2) remains largely unaffected by the different environments.

In addition to altering cation coordination, crowding also affects the hydration of the RNA surface (Figure S3). The number of water molecules within 3.5 nm of the RNA decreases by 14% in the PEG-crowded environment and by a slightly greater 16% under protein crowding.

To better understand the variations in ion binding, we examined cation distributions using radial distribution functions (RDFs) for the three environments (Figure 2C). The RDF profiles from the backbone phosphates, major, and minor grooves indicate that the ion depletion induced by protein crowders primarily occurs non-specifically. While interactions between the RNA binding sites and ions remain consistent between the non-crowded and PEG-crowded conditions, we observed subtle changes in ion distribution within the major and minor grooves of the protein crowders(Figures 2C and S4).

To investigate the mechanism of ion exclusion in protein-crowded environments, we looked at the interactions between Na^+^ and Cl^−^ ions and the crowders (Figure S5). PEG exhibits limited coordination with Na^+^, consistent with its inert and neutral character. In contrast, protein crowders show enhanced interaction with Na^+^ ions, suggesting a competitive exclusion mechanism whereby protein crowders partially sequester ions away from the RNA. This provides a molecular basis for the reduced counterion condensation observed in the protein-crowded condition. This suggests that free cation concentration is lower in protein-crowded conditions, which restricts Na^+^ ion exchange to a greater extent in protein crowders.

Interestingly, despite the significant decrease in Na^+^ ion coordination around the RNA with protein crowders, the electrostatic environment, as measured by the electric flux (Φ_*E*_) (Figure 2D), shows no significant changes at distances greater than 6 Å from the RNA surface across all three conditions. We attribute this partly to the fact that water polarization is the primary contributor to Φ_*E*_, and it is strongest in the non-crowded condition. Water polarization weakens in the presence of crowders, with PEG crowders disrupting water order the most. Notably, the effects of ions and crowder charges appear to compensate each other, resulting in only minor variations in the total Φ_*E*_ at different distances from the RNA surface (Figures S6). As shown in Figure S5, this compensation emerges from distinct contributions of ions, water, and crowder charges to the local electric field. Consequently, the electrostatic potential around the RNA’s salvation shell is minimally affected at the nearest surface (*d* ≈ 0.3 nm). However, at moderate distances (*d* >∼ 0.6 nm), a noticeable reduction in total electrostatic interactions is observed at far distances due to the decreased cation coordination in the protein-crowded environment (Figure 2E).

### Protein Crowders Engage in Chemically Specific RNA Contacts

To further understand RNA interactions in the two crowded conditions, we analyzed pairwise interactions between the RNA and the crowders (Figure 3A). We computed the surface radial distribution function (SRDF), *S*_*X*−*Y*_ (*r*), for different RNA sites in the presence of crowder atoms. Here, *X* represents any atom of the crowding molecule, while *Y* corresponds to the binding surface on RNA, phosphate backbone (PB), major groove (MaG), or minor groove (MiG) of the RNA.

**Figure 3:**
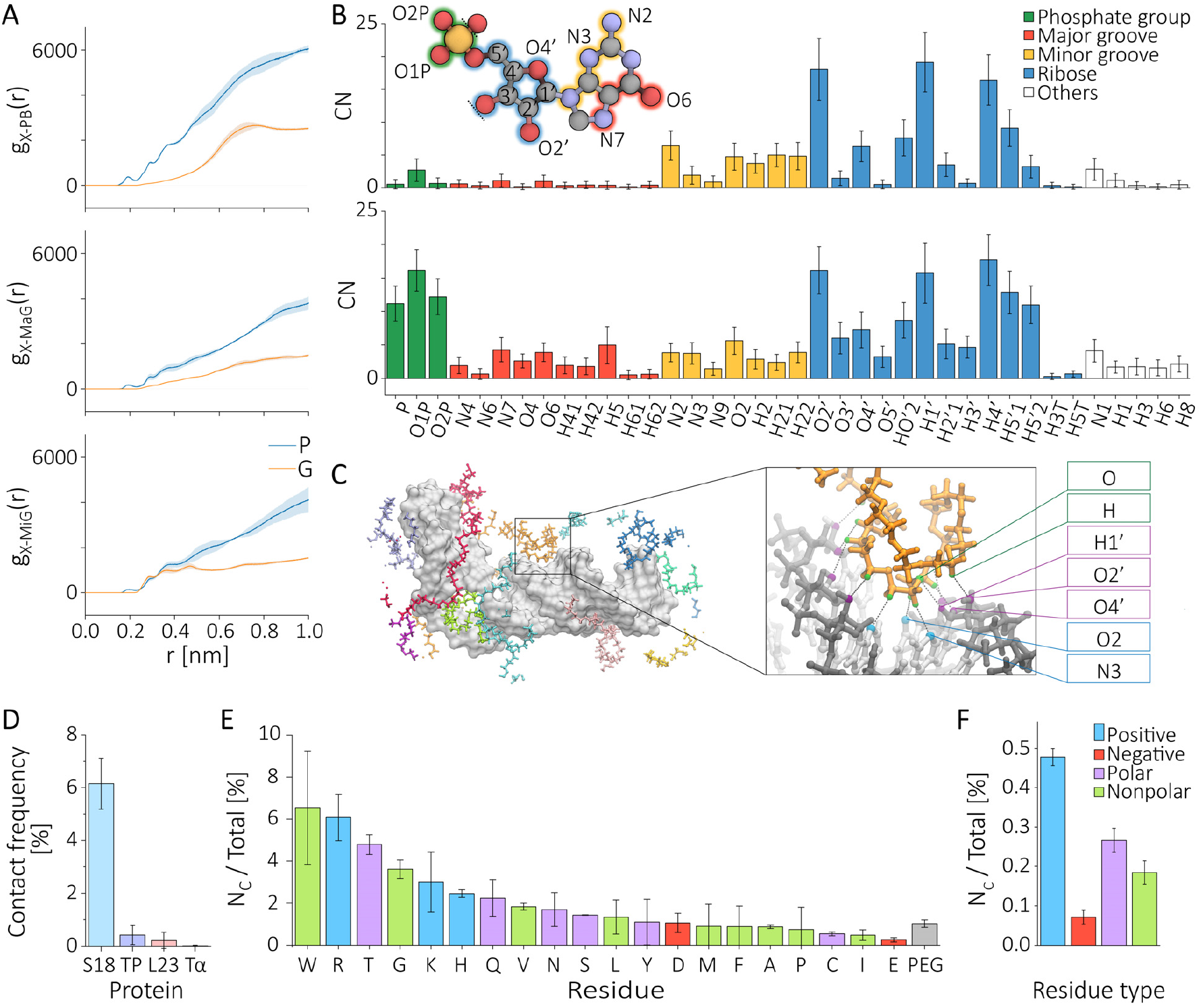
PEG and protein interactions with RNA. A, Surface RDF of PEG (orange) and protein crowders (blue) with the phosphate backbone (*g*_*X*−*PB*_(*r*)), major groove (*g*_*X*−*Mag*_(*r*)), and minor groove (*g*_*X*−*MiG*_(*r*)) of RNA. B, Binding pattern of PEG (top) and proteins (bottom) to RNA within 3.5 Å from the RNA surface. Different regions are visualized along a guanine ribonucleotide in the inset. C, Representation of diverse PEG molecule binding along the RNA, color-coded individually. The RNA is depicted in white. A close-up of the orange PEG highlights its minor groove alignment in the RNA stem, forming hydrogen bonds. The sugar-phosphate backbone (dark gray) and bases (light grey) are shown with interaction sites in purple or blue, respectively. Interacting PEG atoms are highlighted in green. D, Contact frequency of ribosomal protein S18 (S18), translationally controlled tumor protein (TP), ribosomal protein 123 (123), and tubulin alpha lb (T*α*) to the RNA. Cut-off: 8 Å. E-F, Normalized binding frequency of amino acids (E) or residue types (F) to RNA for a 6 Å cutoff. Normalization in E: The number of each residue (name) around the RNA (Figure S7) was divided by the total number of that residue in the simulation box. Normalization in F: The number of residues of the respective type around the RNA was divided by the total number of residues of that type in the simulation box. These detailed contact profiles underscore potential biological relevance, indicating residue-specific interactions that could modulate RNA function and stability in cells.

The SRDF analysis in Figure 3A shows that crowders interact with RNA at both direct (*r* <∼ 3 Å) and indirect (∼ 3 < *r* ∼ 6 Å) distances. Protein crowders coordinate with all defined RNA binding sites, whereas PEG crowders primarily make short-range contacts, mainly to the minor groove. For protein crowders, interactions are mainly with the phosphate backbone atoms and ribose hydroxyl groups (O2^′^, H1^′^, H4^′^, H5^′^1, and H5^′^2), as illustrated in Figure 3B (bottom). In contrast, PEG crowders primarily coordinate with ribose sugars (O2^′^, O4^′^, H1^′^, H4^′^, and HO2^′^) and occasionally with the bases N3 and O2 (Figure 3B, top; Figure 3C). Notably, some PEG molecules bridge the two RNA strands by binding to the minor groove and sugar atoms O2^′^, H1^′^, and O4^′^ (Figure 3C), potentially forming hydrogen bonds with base atoms. Additionally, PEG molecules also bind to exposed bases at hairpin termini, between bulges, and within loop regions.

Unlike the uniform composition of PEG crowders, protein crowders comprise various proteins with different amino acid monomers. To better characterize RNA-protein interactions, we calculated the average contact frequency (Figure 3D), defined as the average probability of a protein being within 8 Å of the RNA. The average is computed by counting that specific protein interaction divided by the protein copies. To account for varying protein lengths, we further normalized the contact frequency by the number of residues in each protein type.

Our analysis reveals that the ribosomal protein S18 (net charge +20) contributes most, occupying a large portion of the RNA surface. Conversely, the negatively charged translationally controlled tumor protein (net charge −11) and tubulin alpha lb (T*α*, net charge −24) rarely contact the RNA. Interestingly, another positively charged ribosomal protein, 123 (net charge: + 17), shows only minor interactions. The lower contact probability of 123, despite its net charge, likely reflects limited accessible surface area or charge burial, emphasizing that net charge alone is not predictive of RNA binding propensity.

We further classified the protein binding in the crowded environment by their constituent amino acid types. To quantify the interaction of individual amino acids, we computed the contact frequency for each residue, normalized by its abundance across all crowders. Shown in (Figure 3F) are our results. Residues directly contacting the RNA are predominantly positively charged, although polar and non-polar residues also contribute significantly. Our contact occupancy analysis indicates a significantly higher contact frequency for proteins compared to PEG crowders (gray bar). Tryptophan, arginine, and threonine are the most frequent contributors to the RNA interface (Figure 3D). We found that tryptophan primarily interacts with exposed bases and sugar atoms, such as those of the 3-nucleotide bulge. Arginine, on the other hand, binds to the sugar-phosphate backbone, particularly around the 3-nucleotide bulge. Threonine, on the other hand, interacts with the RNA’s ribose and single-stranded base atoms, in addition to the 3-nucleotide bulge. These residue-specific interactions not only compete with ions but could potentially influence RNA-protein regulatory interactions in vivo.

### Distinct Crowding Landscapes: PEG Homogeneity vs. Protein Clustering

To compare the two crowding conditions, we analyzed the number density of crowders around the RNA. This involved computing the average density, normalized by the bulk crowder density (*ρ / ρ* _bulk_), for each environment. To visualize the spatial distributions, we took slices along the RNA’s long axis (*z*-axis), as shown in Figure 4A. These slices were then used to generate density maps on the *x-y* plane at various *z* values spanning the RNA’s length. Shown in Figure 4A,B are the number density along the long axis of the RNA.

**Figure 4:**
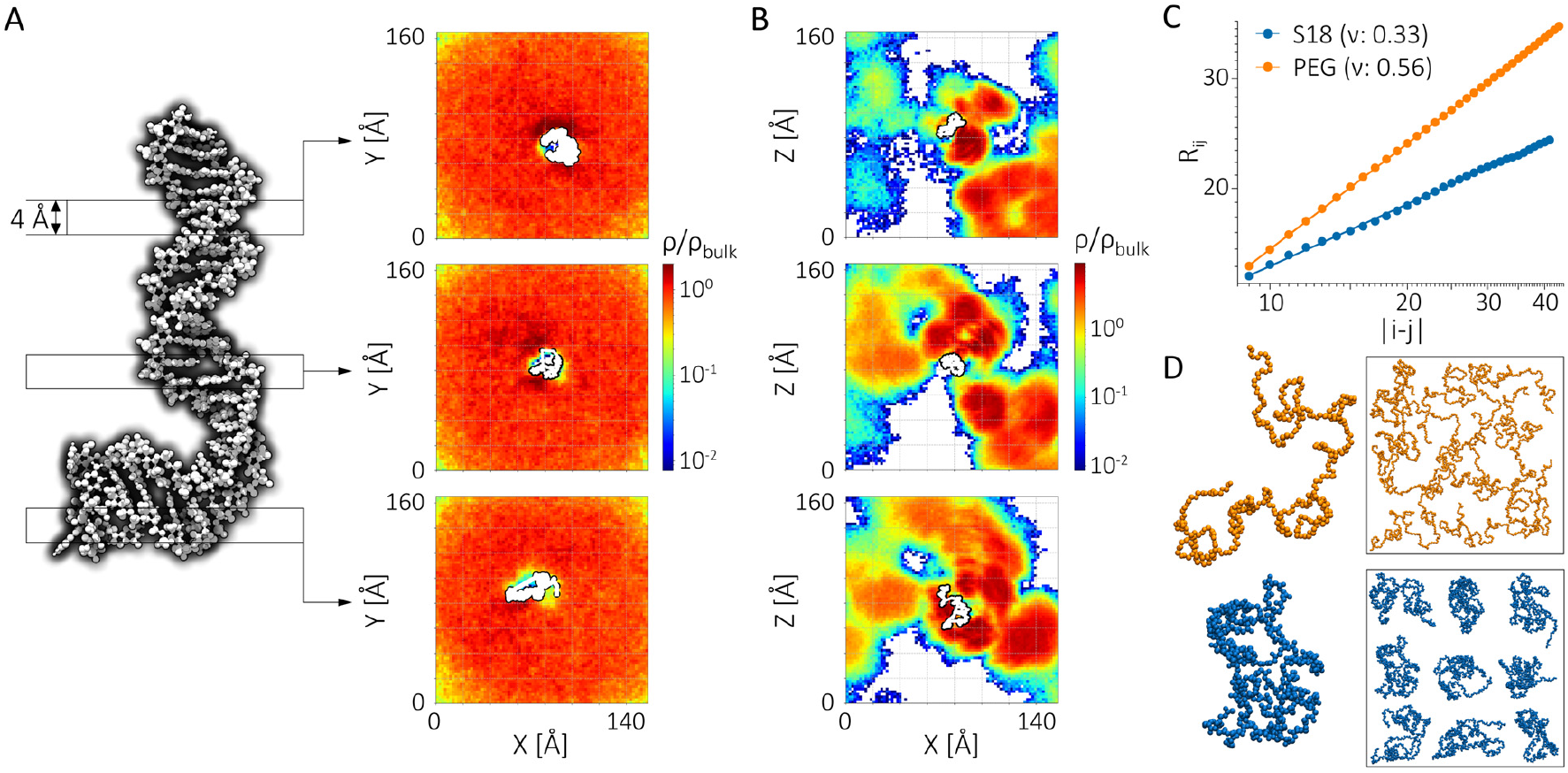
Crowder density distribution around the RNA hairpin. A, PEG- and B, protein-density distribution is given along the x-y coordinates for three 0.4-nm-thick slices of the simulation box taken along the z-axis of the RNA. The RNA is highlighted in white at the center of each frame. C. Flory’s scaling law analysis for a PEG (orange) molecule and a representative S18 protein. The corresponding Flory scaling parameter, *v*, for each crowder is given at the top of the plot. D. Schematic representation of a box with good solvent molecules (*v* > 0.5) and poor solvent molecules (*v* < 0.5), as highlighted here for PEG (orange, top) and S18 molecules (blue, bottom), respectively. PEG *v* ≈ 0.56 confirms good solvent behavior, while S18 *v* ≈ 0.33 indicates clustering due to poor solvent conditions. The distinct Flory exponents illustrate how protein and PEG crowders differently influence RNA microenvironment structure, with proteins creating heterogeneous and clustered environments.

Although PEG molecules are frequently used as crowding agents in experiments ^3,5,39^, they do not necessarily exert identical influences as protein crowders. In our simulations, notable differences emerged in the crowder density distributions between the two environments. PEG molecules displayed a homogeneous distribution around the RNA, with only minor variations near the hairpin region. At this volume fraction, the PEG molecules evenly occupied the available space, indicating that the PEG-RNA system operates in a good-solvent, semi-dilute regime (Figure 4C,D).

In contrast, protein crowders exhibited a markedly heterogeneous distribution, characterized by clusters of high- and low-density regions surrounding the RNA (Figure S8). Note that the average number density of the two crowding states is the same in this study. The distribution of proteins is visualized in Figure S7, which shows crowder clustering and anisotropy around the RNA. We also noted that the average crowder density fluctuated significantly across the slices over the simulation timescale. This behavior arises because water acts as a poor solvent for globular proteins, placing the RNA in a protein environment within a poor-solvent, semi-dilute regime (Figure 4C,D). Flory exponents confirm that PEG behaves as a flexible polymer in good solvent (*v* ≈ 0.56), whereas proteins behave as globular solutes in poor solvent (*v* ≈ 0.33), consistent with crowding-induced clustering.

### Crowder-Dependent Structural Changes in HIV-TAR RNA

As the two crowding conditions create distinct environments, we investigated how variations in crowder type influence the structural ensemble of RNA. To quantitatively assess these differences, we first examined the RNA’s global size using the radius of gyration (*R*_*g*_) and analyzed the number of close contacts between the RNA and crowder atoms (*N*_*c*_). Figure 5A shows these results for the crowders.

**Figure 5:**
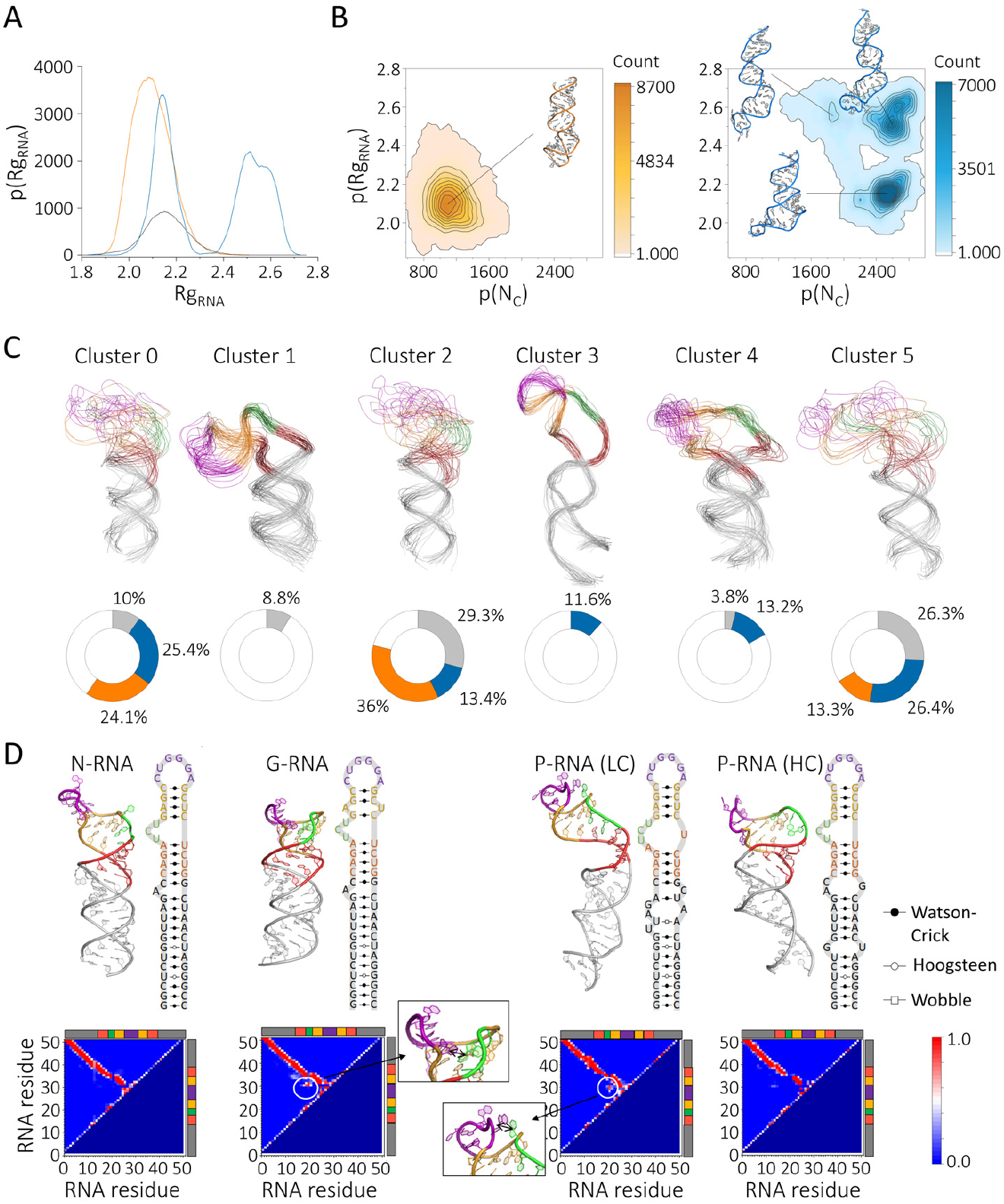
Impact of the environment on RNA structure. A, Histogram of the radius of gyration of the RNA hairpin 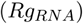 in a non-crowded (grey), PEG-crowded (orange), or protein-crowded (blue) environment. B, Contour representation of 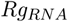 and the number of contacts formed between the RNA and the crowder (PEG: left, highlighted in orange; protein: right, highlighted in blue). C, Representation of the structural population and their distribution within different *in vitro* conditions. Presence of each cluster within the non-crowded, PEG-crowded, or protein-crowded environment is highlighted in the pie chart in grey, orange, and blue, respectively. D, Intramolecular RNA hydrogen bond formation in the different environments. Secondary and tertiary structure formation of the most dominant conformation of the non- (N), PEG- (G), and protein-crowded environments (P). Results for the high (HC) and low crowded (LC) conditions are shown separately. Tertiary structure formation is given as the number of contacts formed between the residues throughout the simulation trajectory. The inset shows the structural changes that led to new residue contact formations. The apical loop, upper stem, 3-nt bulge, lower stem, and extended stem are highlighted in purple, yellow, green, red, and grey, respectively.

Uniform PEG crowders resulted in approximately *N*_*c*_ ≈ 1100 contacts, with the RNA slightly compressed to an average *R*_*g*_ of 2.1 nm, compared to 2.16 nm under non-crowded conditions (Figure 5B). In contrast, protein crowders formed nearly twice as many contacts as PEG crowders at the same concentration (Figures 5B and S8). Interestingly, RNA expansion was observed at higher crowder densities (high *N*_*c*_). Our observation of RNA expansion under protein crowding is consistent with Ref^65^, who predicted that rigid DNA structures could undergo expansion or compaction depending on crowder type and interactions, suggesting a nuanced interplay between steric and electrostatic factors. Overall, PEG crowders tend to compress the RNA, whereas protein mixtures, which mimic a cellular environment, can induce RNA expansion.

Beyond global size changes, RNA-crowder interactions also influenced local structural features. Specifically, protein crowders caused the major grooves to widen about 40% (Figure S9) and as a result the minor grooves to narrow. Groove remodeling may alter accessibility to RNA-binding proteins, small molecules, or ribonucleoprotein partners, especially at sites where major groove recognition plays a functional role. While groove dimensions were minimally impacted by PEG crowders.

These changes in RNA size and groove dimensions resulted in conformational ensembles that varied with the environment, as illustrated in Figure 5C. In PEG-crowded environments, RNA conformational state clusters around 0, 2, and 5, which were characterized as a high hairpin disordered state, where the upper hairpin region (residues 19-35) is disorganized. In contrast, RNA in protein-crowded environments explored a mix of ordered and disordered loops, with wider major grooves and A-form groove distances, leading to more diverse conformational states. The ensembles sampled by RNA in protein crowders and in a dilute regime show similarity. Yet there are still unique conformational states exclusive to protein-crowded conditions (cluster 3) and non-crowded conditions (cluster 1). These results suggest that the conformational ensemble of HIV-1 TAR RNA is modulated by its surrounding environment.

To examine secondary and tertiary contact changes induced by different environments, we selected the most populated cluster from each simulation setup and constructed its secondary and tertiary contact maps (Figure 5D, left). In both non-crowded and PEG-crowded envi-ronments, tertiary contacts primarily formed in the upper 4-bp stem region, with additional contacts observed between the apical loop and the 3-nt bulge in PEG-crowded conditions. In contrast, RNA in protein-crowded environments showed notable shifts in both secondary and tertiary contacts (Figure 5D, right). Specifically, the lower hairpin region, comprising the 5-bp and 12-bp helices and the 1-nt bulge, exhibited major alterations in intramolecular hydrogen bond formation. High-density protein-crowding conditions led to loss of hydrogen bonds, while low protein-crowding conditions induced altered hydrogen bond formation around the bulge region. Additionally, in high protein-crowding environments, the 3-nt bulge is separated from the upper 4-bp stem, resulting in loss of contacts.

We also investigated how the RNA’s environment influences its structural dynamics. Supplementary Movie S1 visually compares the RNA’s structural behavior across the three environments. To quantify structural dynamics, we analyzed backbone flexibility by computing root mean square fluctuations (RMSF) that relate to the experimentally available B-factors. We also estimate SHAPE reactivities from MD trajectories, as described in previous studies ^63,64^ and detailed in the methods section. The results of these analyses are shown in Figure 6.

**Figure 6:**
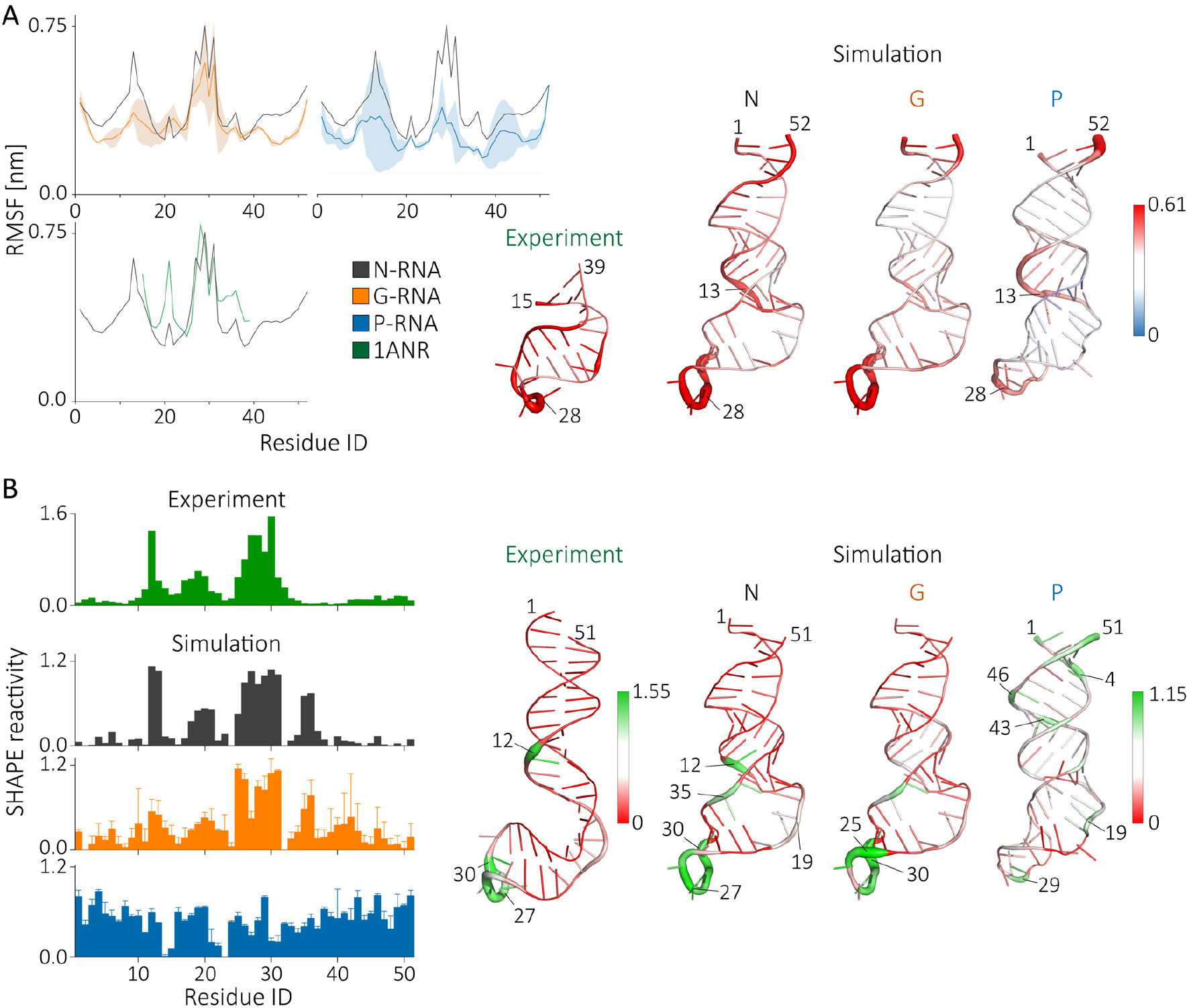
Effects of the environment on RNA residue and backbone flexibility. A. Root-mean-square fluctuation (RMSF) per residue and the corresponding B-factor representation for the experimental-determined RNA structure (green), and the simulation of the RNA in non-crowded (N, gray), PEG-crowded (G, orange), and protein-crowded (P, blue) environments. Experimental values were calculated based on the twenty conformers comprising the 1ANR PDB entry, obtained in the presence of 50 mM NaCl ^67^. Data for the PEG- and protein-rich solutions are presented as average values calculated from two different simulations, with the standard deviation indicated by light orange and blue shading, respectively. B, Experimental (E, green) and calculated RNA backbone flexibility per residue with corresponding structural representation. Experimental data is taken from reference ^66^.

First, we assessed how well RMSF calculations reflect the B-factor data of the HIV TAR measured in a dilute regime. The calculated RMSF is consistent with experimental data (Figure 6A bottom). The agreement between simulation and experiment in the dilute state encouraged us to compare the dynamics in crowding conditions. RMSF analysis revealed minimal differences between non-crowded and PEG-crowded environments (Figure 6A). In contrast, RNA in protein-crowded environments exhibited suppressed flexibility in the apical loop (residues 26-31), the 1-nt bulge (residue 13), and the 3-nt bulge (residues 19-21) (Figure 6A, right).

To explore how crowding affects SHAPE reactivities, we first validated our computational methodology using *in vitro* SHAPE reactivity data of HIV-1 TAR RNA ^66^ (Figure 6B, green-black) in dilute conditions. The strong agreement between experimental and simulation profiles supports the validity of our approach, enabling comparisons of SHAPE reactivities derived from simulations under crowding conditions where experimental data are unavailable.

Consistent with *in vitro* experiments, increased backbone flexibility was detected in the bulge and apical loop regions. The simulation data also showed a weaker peak on the opposite strand of the 3-nt bulge. RNA behavior in PEG-crowded environments closely mirrored *in vitro* SHAPE reactivities (Figure 6B, orange), with reduced reactivity in 12-15 nt while major changes are observed at the protein-crowded environments (Figure 6B, blue vs black). Overall, we observed a pronounced reduction in computed SHAPE reactivities in protein crowders. Highly mobile regions of the RNA (residues 12-16 and 25-32) are dramatically suppressed due to specific protein binding. We observe a minor increase in relative reactivity at the stem region, likely due to an expanded major groove unique to the protein crowders. These findings underscore the distinct effects of protein-crowding environments on the structure and dynamics of RNA and, more importantly, suggest that SHAPE reactivity in crowding conditions does not report on the dynamics as it does in vitro.

### RNA and Solvent Dynamics Depend Critically on Crowder Identity

The results presented so far demonstrate how crowders affect RNA structure and the surrounding water-cation environment. Here, we delve into the specific consequences of the crowding environment on RNA and solvent dynamics, including translational and rotational diffusion, dynamic structural responses, and the relaxation of the solvation shell. Details of computing these properties are described in the methods section.

Figure 7A compares the diffusion constants of water molecules, Na^+^ ions, and RNA, highlighting the impact of crowding on lateral mobility. Crowding consistently reduces lateral diffusion for all components. Relative to the dilute environment, water diffusion decreases by 42% in PEG and by 26% in protein-crowded conditions. The diffusion of Na^+^ ions is impeded even more significantly, decreasing by 66% with PEG and by an average of 43% with proteins. Notably, the RNA experiences the most substantial slowdown; the dense protein environment drastically reduces RNA diffusion by approximately 37-fold (from 158 to 5 *µ*m^2^ /s). In contrast, PEG crowders induce a comparatively modest 8-fold reduction (from 158 to 15 *µ*m^2^/s).

**Figure 7:**
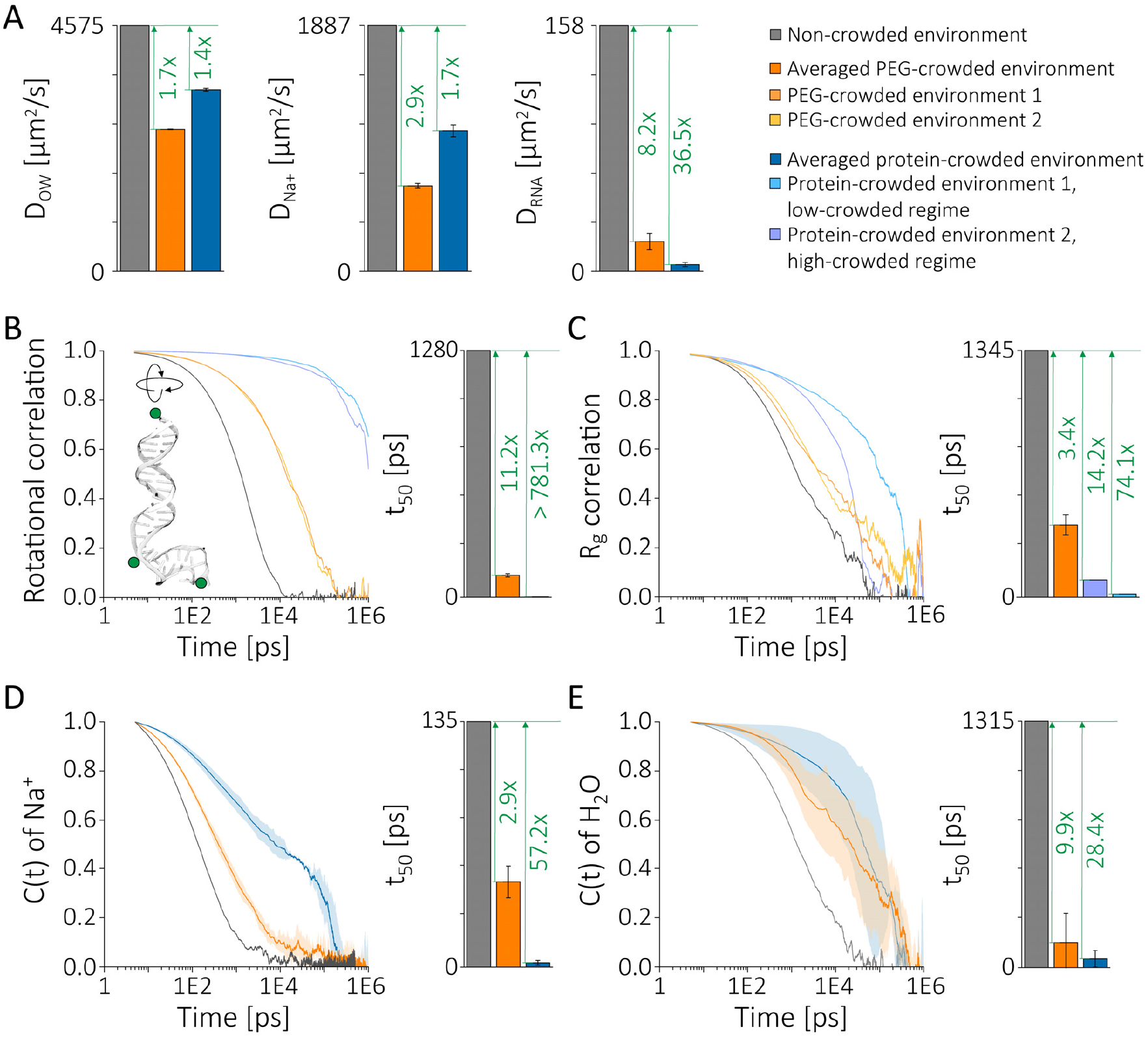
Influence of crowding environments on RNA and solvent dynamics. (A) Diffusion coefficients highlighting dramatic RNA translational slowdown under protein crowding compared to PEG conditions. (B) Rotational correlation times, showing substantial decoupling from translational diffusion under protein crowding. (C) Structural relaxation dynamics of RNA radius of gyration (*R*_*g*_ ). (D, E) Time correlation analysis of surface-bound Na^+^ ions and water molecules. These results demonstrate that protein crowders uniquely and significantly slow down RNA and local solvent dynamics, emphasizing crowder identity as a crucial determinant of biomolecular dynamics.

Crowding similarly exerts a profound influence on RNA rotational dynamics (Figure 7B). The rotational correlation times indicate roughly an 800-fold slowdown in RNA tumbling under protein crowding, suggesting a clear decoupling of translational and rotational dynamics due to distinctly separated timescales. In contrast, PEG crowders have a smaller effect on RNA rotation, maintaining closer coupling between translational and rotational degrees of freedom.

Next, we investigated internal RNA dynamics through the autocorrelation of the radius of gyration *R*_*g*_ (Figure 7C). RNA conformational dynamics in the protein-crowded environment slowed significantly (by approximately 14-74-fold) compared to the dilute solution. PEG crowding, however, resulted in a relatively modest 60% slowdown.

A striking difference emerges when comparing solvent and RNA dynamics: bulk mobility of ions and water is more strongly affected by PEG than protein crowders, while RNA dynamics are much more drastically reduced under protein crowding. To clarify this contrast, we analyzed the dynamics of counterions and water specifically at the RNA surface, as surface solvent dynamics differ notably from bulk behaviors.

Figure 7D shows time-dependent correlations for Na^+^ ions within 7Å (Debye length) and water molecules within 3.5 Å of the RNA surface. Protein crowding markedly slowed counterion dynamics by approximately 57-fold and water dynamics by around 28-fold. In contrast, PEG crowders slowed these dynamics by only 3-10 fold. This sharp contrast between surface and bulk solvent dynamics highlights that PEG primarily affects bulk solvent mobility (Figure 7A), whereas protein crowders predominantly impact RNA surface dynamics.

These marked differences between local and bulk solvent dynamics under protein and PEG crowding indicate that protein-crowder interactions specifically and selectively modulate RNA surface hydration and ion mobility, fundamentally altering the solvent environment around RNA relative to bulk behavior.

Overall, our simulations reveal substantial slowdowns in solvent and RNA internal motions under crowded conditions. Specifically, we observed RNA internal dynamics slowing by approximately 14-75 fold, water mobility by about 30-fold, and Na^+^ ion dynamics by roughly 60-fold. These values are consistent with or exceed experimental measurements reported for comparable systems. For instance, yeast tRNAs show approximately an 8-fold reduction in cytoplasmic diffusion under mild crowding, reaching up to an 80-fold slowdown under severe osmotic stress conditions ^68^. Similarly, DNA diffusion typically slows 5-20 fold in polymer-crowded solutions, depending on crowder characteristics and ionic conditions ^69^, closely matching our PEG-crowded simulations. Additionally, studies report significant slow-downs in rotational than translational dynamics for proteins ^70^, aligning qualitatively with our observations. Although direct experimental measurements of ion dynamics under crowding remain limited, theoretical studies predict substantial hindrance due to protein-crowder electrostatic interactions^27,71^, corroborating our simulation results. These comparisons validate our molecular dynamics simulations as realistic models for understanding crowding effects in biological environments and underscore the generalizability of these crowding-induced dynamics across different biomolecular systems.

## Discussion

Understanding how macromolecular crowding reshapes the structural and electrostatic landscape of RNA is essential for translating in vitro biophysical insights into the context of the living cell. Despite advances in theory and modeling, atomistic-level characterization of RNA-ion-crowder interactions under physiologically relevant conditions remains scarce. Our study addresses this gap by using microseconds-long, all-atom molecular dynamics simulations to directly compare the impact of synthetic (PEG) and biological (protein) crowders on the hydration, ion coordination, and conformational dynamics of the HIV-1 TAR RNA hairpin. By revealing how protein identity, not just crowding concentration, modulates RNA behavior at the molecular level, this work informs both RNA-based therapeutics and in vivo structure-function modeling. Our atomistic simulations are the first, to our knowledge, to explicitly demonstrate at a molecular level how specific electrostatic and hydration interactions under protein crowding environments fundamentally reshape RNA structural ensembles and dynamics. This contrasts starkly with PEG-based crowding studies that overlook these critical physicochemical details.

Our all-atom simulations revealed divergent effects of PEG and protein crowding on RNA, particularly concerning its electrostatic environment and hydration shell. We observed that PEG crowding provides an inadequate mimic of the protein-rich cellular environment, more closely resembling non-crowded *in vitro* conditions. This discrepancy arises from PEG’s lack of diverse intermolecular interactions compared to the heterogeneous protein environment. Our findings align with the concept of a transient intracellular environment, where crowding fluctuates due to protein interactions ^72^. The distinct distributions of PEG and protein crowders significantly impact RNA structure and the dynamics of the RNA and surrounding solvent.

While the global fold of the HIV-TAR hairpin remained stable across all conditions, we observed distinct structural variations. The heterogeneous protein crowding generated a more diverse structural population, including unique clusters not seen under other conditions. This observation is consistent with experimental findings indicating that certain *in vivo* RNA conformations cannot be replicated *in vitro* ^11,23^. Conversely, PEG primarily induced compaction, a general consequence of excluded volume favoring more compact biomolecular states^10,18^. Notably, our results hint that changes in experimental SHAPE signals could reflect backbone distortion due to protein interactions rather than RNA denaturation ^73^.

In contrast to the structural effects, the electrostatic environment, as assessed by *Na^+^* condensation, remained largely unaffected by PEG, as in dilute solutions. This sharply contrasts with the protein-crowded environment, where a key finding was a substantial ( ∼40%) reduction in *Na^+^* counterion condensation around the RNA. This reduction in cation condensation suggests that protein crowders significantly reshape the RNA’s electrostatic landscape. This aligns with predictions from spatially heterogeneous dielectric decrement theory ^71,74^, in which ion depletion arises from competition with charged protein surfaces and reduced solvent polarizability. Our data validate these models and provide the first direct atomistic evidence of ion exclusion via protein-RNA contact. This reduction arises from several non-specific interactions between protein crowders and the RNA surface, particularly involving positively charged residues like arginine. This is consistent with previous studies highlighting the importance of arginine in RNA-protein interactions^12,75^. Although our results are based on HIV-TAR, known to form specific interactions with arginine in the TAR:Tat RBD complex46-50, we anticipate similar non-specific interactions with other RNA structures. These interactions likely displace *Na^+^* ions from their typical binding sites along the RNA backbone and within the major groove, emphasizing that biological crowders influence RNA behavior not only through volume exclusion but also by competing for ion binding sites, thereby altering the electrostatic potential experienced by the RNA.

A particularly significant finding from our work is the dramatic slowdown of RNA translational and rotational dynamics under protein-crowded conditions, which far exceeds the effects observed in PEG solutions. Our observed slowdown factors–approximately 37-fold for translational diffusion and up to 800-fold for rotational tumbling–align qualitatively with previous experimental and theoretical studies. Yeast tRNAs exhibit similar, although slightly smaller, slowdowns (approximately 8–80–fold) in cytoplasmic conditions depending on the level of osmotic stress^68^. Similarly, DNA in polymer solutions slows by around 5–20-fold ^69^, reflecting a broader trend where crowding-induced slowdowns depend critically on crowder identity and physicochemical interactions. Our water and ion dynamics slowdowns (approximately 30- and 60-fold, respectively) further confirm theoretical predictions that crowding strongly affects hydration layer dynamics and ion mobility ^71^. These results emphasize the crucial need to accurately represent crowder interactions in both simulation and experimental studies to faithfully capture biomolecular dynamics in cellular environments.

The differential impact on ion condensation has implications for RNA stability. While crowding generally enhances RNA stability through excluded volume effects, the protein-induced reduction in counterion condensation could potentially destabilize the RNA by weakening crucial electrostatic stabilization ^25,35^. However, the maintained global fold and the indication of higher electrostatic stability in our simulations suggest that direct protein-RNA interactions compensate for this loss of ionic stabilization, highlighting the complexity of biological crowding compared to synthetic crowding.

Intriguingly, our simulations also revealed non-specific interactions between PEG and the RNA, specifically with ribose atoms along the minor groove and with nucleobase atoms in single-stranded regions. These interactions involving the sugar moiety align with previous reports of reduced β-galactosidase activity, potentially due to PEG’s interference with galactose substrate binding ^76^. While these are non-specific, they may have implications for PEG-RNA conjugate design, drug delivery, or shielding effects in therapeutic contexts. The presence of these PEG-RNA interactions suggests that a purely excluded-volume model is insufficient to fully account for the effects of PEG crowding on RNA and may have implications for PEG-based therapeutics ^77^.

Furthermore, our simulations indicate that both types of crowding influence the RNA’s salvation environment. PEG reduces bulk water mobility, consistent with prior findings ^78^, and decreases both the number and dynamic behavior of water molecules surrounding the RNA. In contrast, protein crowders had a lesser impact on bulk water dynamics but a higher impact on reducing the number of surrounding water molecules and altering the RNA-water hydrogen bonding network. This likely arises from direct protein-RNA interactions that modulate hydration, possibly through localized dehydration or changes in water structuring.

The inability of PEG to replicate the electrostatic perturbations induced by protein-rich crowding underscores the limitations of using PEG as a universal model for cellular crowding, particularly when investigating phenomena sensitive to electrostatics and hydration, both critical for RNA function. Our findings support studies suggesting that synthetic crowders may not fully capture the intricate interplay of forces present in the cell. ^11,33^

## Summary

In conclusion, our atomic-level simulations demonstrate that protein and PEG crowders exert distinct influences on RNA electrostatics and hydration. Protein crowding significantly reduces counterion condensation through direct interactions, likely altering the RNA’s hydration landscape, whereas PEG primarily acts through volume exclusion with minimal impact on the ionic environment. These findings underscore the need for caution when using synthetic crowders to mimic the complex effects of the cellular environment on RNA. Consequently, exploring more biologically relevant crowding agents is essential for a more accurate understanding of RNA behavior *in vivo*.

Future studies integrating MD with SAXS, SHAPE-MaP, or ion-sensitive FRET could test the predictions here, including hydration loss and counterion displacement under crowding. Our findings are likely generalizable to other structured RNAs such as riboswitches, ribozymes, or long non-coding RNAs, where electrostatics and hydration play similarly critical roles. The structural features impacted here, major/minor groove geometry, local hydration, and cation binding, are not unique to TAR RNA but are conserved across many structured RNAs. This suggests that riboswitches, ribozymes, and long non-coding RNAs may similarly experience altered electrostatic landscapes under protein-rich crowding, affecting folding, ligand binding, or catalysis. Thus, the ion-exclusion and hydration-remodeling effects observed here are likely to influence folding and recognition in riboswitches, pseudoknots, and viral replication elements. Our predictions could be tested using site-specific SHAPE reactivity assays or osmotic stress experiments in PEG vs. protein-crowded solutions, focusing on bulge and groove-associated reactivity changes. Additionally, time-resolved FRET or neutron scattering could validate hydration dynamics at nanosecond timescales. Such validation would provide critical insight into ion competition and local hydration dynamics at biologically relevant RNA motifs, further bridging the gap between computational predictions and biological reality.

## Acknowledgement

This work was supported by the NYUAD Faculty support grant AD181 to SK. Computer simulations were carried out on the High-Performance Computing resources at New York University Abu Dhabi.

## References

(1) Zimmerman, S. B.; Trach, S. O. Estimation of macromolecule concentrations and excluded volume effects for the cytoplasm of Escherichia coli. Journal of Molecular Biology 1991, 222, 599–620.

(2) Ellis, R. J. Macromolecular crowding: obvious but underappreciated. Trends in Biochemical Sciences 2001, 26, 597–604.

(3) Rivas, G.; Minton, A. P. Macromolecular crowding in vitro, in vivo, and in between. Trends in Biochemical Sciences 2016, 41, 970–981.

(4) Theillet, F.-X.; Binolfi, A.; Frembgen-Kesner, T.; Hingorani, K.; Sarkar, M.; Kyne, C.; Li, C.; Crowley, P. B.; Gierasch, L.; Pielak, G. J.; Elcock, A. H.; Gershenson, A.; Selenko, P. Physicochemical Properties of Cells and Their Effects on Intrinsically Disordered Proteins (IDPs). Chemical Reviews 2014, 114, 6661–6714.

(5) Zhou, H. X.; Rivas, G.; Minton, A. P. Macromolecular crowding and confinement: Biochemical, biophysical, and potential physiological consequences. Annual Review of Biophysics 2008, 37, 375 397.

(6) Minton, A. P. Implications of macromolecular crowding for protein assembly. Current Opinion in Structural Biology 2000, 10, 34–39.

(7) van den Berg, B.; Ellis, R. J.; Dobson, C. M. Effects of macromolecular crowding on protein folding and aggregation. EMBO Journal 1999, 18, 6927 6933.

(8) Tokuriki, N.; Kinjo, M.; Negi, S.; Hoshino, M.; Goto, Y.; Urabe, I.; Yomo, T. Protein folding by the effects of macromolecular crowding. Protein Science 2004, 13, 125–133.

(9) Cheung, M. S.; Klimov, D.; Thirumalai, D. Molecular crowding enhances native state stability and refolding rates of globular proteins. Proceedings of the National Academy of Sciences 2005, 102, 4753–4758.

(10) Kilburn, D.; Roh, J. EL; Guo, L.; Briber, R. M.; Woodson, S. A. Molecular crowding stabilizes folded RNA structure by the excluded volume effect. Journal of the American Chemical Society 2010, 132, 8690–8696.

(11) Tyrrell, J.; Weeks, K. M.; Pielak, G. J. Challenge of Mimicking the Influences of the Cellular Environment on RNA Structure by PEG-Induced Macromolecular Crowding. Biochemistry 2015, 54, 6447 6453.

(12) Collauto, A.; von Bülow, S.; Gophane, D. B.; Saha, S.; Stelzl, L. S.; Hummer, G.; Sigurdsson, S. T.; Prisner, T. F. Compaction of RNA Duplexes in the Cell. Angewandte Chemie 2020, 59, 23025–23029.

(13) Nakano, S.; Miyoshi, D.; Sugimoto, N. Effects of molecular crowding on the structures, interactions, and functions of nucleic acids. Chemical Reviews 2014, 114, 2733–2758.

(14) Bloomfield, V. A. DNA condensation. Current Opinion in Structural Biology 1996, 6, 334–341.

(15) Zinchenko, A. A.; Yoshikawa, K. Na^+^ Shows a Markedly Higher Potential than K+ in DNA Compaction in a Crowded Environment. Biophysical Journal 2005, 88, 4118–4123.

(16) Hirano, K.; Ichikawa, M.; Ishido, T.; Ishikawa, M.; Baba, Y.; Yoshikawa, K. How environmental solution conditions determine the compaction velocity of single DNA molecules. Nucleic Acids Research 2012, 40, 284–289.

(17) Cheng, C.; Jia, J.-L.; Ran, S.-Y. Polyethylene glycol and divalent salt-induced DNA reentrant condensation revealed by single molecule measurements. Soft Matter 2015, 11, 3927–3935.

(18) Denesyuk, N. A.; Thirumalai, D. Crowding promotes the switch from hairpin to pseu-doknot conformation in human telomerase RNA. Journal of the American Chemical Society 2011, 133, 11858 11861.

(19) Denesyuk, N. A.; Thirumalai, D. Entropic stabilization of the folded states of RNA due to macromolecular crowding. Biophysical Reviews 2013, 5, 225–232.

(20) Kilburn, D.; Roh, J. H.; Behrouzi, R.; Briber, R. M.; Woodson, S. A. Crowders perturb the entropy of RNA energy landscapes to favour folding. Journal of the American Chemical Society 2013, 135, 10055–10063.

(21) Desai, R.; Kilburn, D.; Lee, H.-T.; Woodson, S. A. Increased Ribozyme Activity in Crowded Solutions. Journal of Biological Chemistry 2014, 289, 2972–2977.

(22) Strulson, C. A.; Boyer, J. A.; Whitman, E. E.; Bevilacqua, P. C. Molecular crowders and cosolutes promote folding cooperativity of RNA under physiological ionic conditions. RNA 2014, 20, 331–347.

(23) Tyrrell, J.; McGinnis, J. L.; Weeks, K. M.; Pielak, G. J. The Cellular Environment Stabilizes Adenine Riboswitch RNA Structure. Biochemistry 2013, 52, 8777–8785.

(24) Nakano, S.-i.; Kitagawa, Y.; Yamashita, H.; Miyoshi, D.; Sugimoto, N. Effects of Cosolvents on the Folding and Catalytic Activities of the Hammerhead Ribozyme. ChemBioChem 2015, 16, 1803–1810.

(25) Sorin, E. J.; Rhee, Y. M.; Pande, V. S. Does water play a structural role in the folding of small nucleic acids? Biophysical Journal 2005, 88, 2516–2524.

(26) Lambert, D.; Draper, D. E. Effects of osmolytes on RNA secondary and tertiary structure stabilities and RNA-Mg^2^ interactions. Journal of Molecular Biology 2007, 370, 993-1005.

(27) Manning, G. S. The molecular theory of polyelectrolyte solutions with applications to the electrostatic properties of polynucleotides. Quarterly Reviews of Biophysics 1978, 11, 179 246.

(28) Yu, T.; Zhu, Y.; He, Z.; Chen, S.-J. Predicting Molecular Crowding Effects in lon-RNA Interactions. The Journal of Physical Chemistry B 2016, 120, 8837–8844.

(29) Nakano, M.; Tateishi-Karimata, H.; Tanaka, S.; Tama, F.; Miyashita, O.; Nakano, S.-i.; Sugimoto, N. Thermodynamic properties of water molecules in the presence of cosolute depend on DNA structure: a study using grid inhomogeneous solvation theory. Nucleic Acids Research 2015, 43, 10114–10125.

(30) Harries, D.; Rösgen, J. A practical guide on how osmolytes modulate macromolecular properties. Methods in Cell Biology 2008, 84, 679–735.

(31) Ghosh, S.; Takahashi, S.; Ohyama, T.; Liu, L.; Sugimoto, N. Elucidating the Role of Groove Hydration on Stability and Functions of Biased DNA Duplexes in Cell-Like Chemical Environments. Journal of the American Chemical Society 2024, 146, 32479–32497.

(32) Lambert, D.; Draper, D. E. Effects of osmolytes on RNA−Mg^2^ interactions: implications for the folding of RNA in vivo. Journal of Molecular Biology 2005, 345, 907 917.

(33) Daher, M.; Widom, J. R.; Tay, W.; Walter, N. G. Soft Interactions with Model Crowders and Non-canonical Interactions with Cellular Proteins Stabilize RNA Folding. Journal of Molecular Biology 2018, 430, 509–523.

(34) Holmstrom, E. D.; Liu, Z.; Netteis, D.; Best, R. B.; Schuler, B. Disordered RNA chaperones can enhance nucleic acid folding via local charge screening. Nature Communications 2019, 10, 2453.

(35) Speer, S. L.; Stewart, C. J.; Sapir, L.; Harries, D.; Pielak, G. J. Macromolecular Crowding Is More than Hard-Core Repulsions. Annual Review of Biophysics 2022, 51, 267–300.

(36) Liebau, J.; Laatsch, B. F.; Rusnak, J.; Gunderson, K.; Finke, B.; Bargender, K.; Narkiewicz-Jodko, A.; Weeks, K.; Williams, M. T.; Shulgina, I.; Musier-Forsyth, K.; Bhattacharyya, S.; Hati, S. Polyethylene Glycol Impacts Conformation and Dynamics of Escherichia coli Prolyl-tRNA Synthetase Via Crowding and Confinement Effects. Biochemistry 2024, 63, 1621–1635.

(37) Zohra, F. T.; Al-Zuhairi, H.; Reinoza, J.; Kim, H.; Hanke, A. Probing the effect of PEG-DNA interactions and buffer viscosity on tethered DNA in shear flow. PLOS ONE 2025, 20, e0329961.

(38) Parray, Z. A.; Hassan, M. I.; Ahmad, F.; Islam, A. Amphiphilic nature of polyethylene glycols and their role in medical research. Polymer Testing 2020, 82, 106316.

(39) Miklos, A. C.; Li, C.; Sharaf, N. G.; Pielak, G. J. Volume exclusion and soft interaction effects on protein stability under crowded conditions. Biochemistry 2010, 49, 6984–6991.

(40) Draper, D. E. A guide to ions and RNA structure. RNA 2004, 10, 335 343.

(41) Tan, Z.-J.; Chen, S.-J. Nucleic Acid Helix Stability: Effects of Salt Concentration, Cation Valence and Size, and Chain Length. Biophysical Journal 2006, 90, 1175–1190.

(42) Yoo, J.; Aksimentiev, A. Improved Parameterization of Amine-Carboxylate and Amine-Phosphate Interactions for Molecular Dynamics Simulations Using the CHARMM and AMBER Force Fields. Journal of Chemical Theory and Computation 2016, 12, 430–443.

(43) Zgarbová, M.; Otyepka, M.; Šponer, J.; Mládek, A.; Banáš, P.; Cheatham, T. E.; Jurecka, P. Refinement of the Cornell et al. Nucleic Acids Force Field Based on Reference Quantum Chemical Calculations of Glycosidic Torsion Profiles. Journal of Chemical Theory and Computation 2011, 7, 2886 2902.

(44) Joung, I. S.; Cheatham, T. E. Determination of Alkali and Halide Monovalent Ion Parameters for Use in Explicitly Solvated Biomolecular Simulations. Journal of Physical Chemistry B 2008, 112, 9020–9041.

(45) Jorgensen, W. L.; Chandrasekhar, J.; Madura, J. D.; Impey, R. W.; Klein, M. L. Comparison of simple potential functions for simulating liquid water. The Journal of Chemical Physics 1983, 79, 926–935.

(46) Puglisi, J. D.; Chen, L.; Blanchard, S.; Frankel, A. D. Solution structure of a bovine immunodeficiency virus Tat-TAR peptide-RNA complex. Science 1995, 270, 1200–1203.

(47) Puglisi, J. D.; Chen, L.; Frankel, A. D.; Williamson, J. R. Role of RNA structure in arginine recognition of TAR RNA. Proceedings of the National Academy of Sciences 1993, 90, 3680–3684.

(48) Puglisi, J. D.; Tan, R.; Calnan, B. J.; Frankel, A. D.; Williamson, J. R. Conformation of the TAR RNA-Arginine Complex by NMR Spectroscopy. Science 1992, 257, 76 80.

(49) Tao, J.; Chen, L.; Frankel, A. D. Dissection of the Proposed Base Triple in Human Immunodeficiency Virus TAR RNA Indicates the Importance of the Hoogsteen Interaction. Biochemistry 1997, 36, 3491–3495.

(50) Ken, M. L.; Roy, R.; Geng, A.; Ganser, L. R.; Manghrani, A.; Cullen, B. R.; Schulze-Gahmen, U.; Herschlag, D.; Al-Hashimi, H. M. RNA conformational propensities determine cellular activity. Nature 2023, 617, 835–841.

(51) Henning-Knechtel, A.; Thirumalai, D.; Kirmizialtin, S. Differences in ion-RNA binding modes due to charge density variations explain the stability of RNA in monovalent salts. Science Advances 2022, 8, eabo1190.

(52) Freund, J.; Kalbitzer, H. R. Physiological buffers for NMR spectroscopy. Journal of Biomolecular NMR 1995, 5, 321–322.

(53) Misra, V. K.; Draper, D. E. A thermodynamic framework for Mg2+ binding to RNA. Proceedings of the National Academy of Sciences 2001, 98, 12456–12461.

(54) Feig, M.; Yu, I.; Wang, P.-h.; Nawrocki, G.; Sugita, Y. Crowding in Cellular Environments at an Atomistic Level from Computer Simulations. The Journal of Physical Chemistry B 2017, 121, 8009–8025.

(55) The Human Protein Atlas. 2023; https://www.proteinatlas.org.

(56) Miyamoto, S.; Kollman, P. A. Settle: An analytical version of the SHAKE and RATTLE algorithm for rigid water models. Journal of Computational Chemistry 1992, 13, 952–962.

(57) Hess, B.; Bekker, H.; Berendsen, H. J. C.; Fraaije, J. G. E. M. LINCS: A linear constraint solver for molecular simulations. Journal of Computational Chemistry 1997, 18, 1463-1472.

(58) Bussi, G.; Donadio, D.; Parrinello, M. Canonical sampling through velocity rescaling. The Journal of Chemical Physics 2007, 126, 014101.

(59) Parrinello, M.; Rahman, A. Polymorphic transitions in single crystals: A new molecular dynamics method. Journal of Applied Physics 1981, 52, 7182 7190.

(60) Ng, A. Y.; Jordan, M. L; Weiss, Y. On Spectral Clustering: Analysis and an Algorithm. Advances in Neural Information Processing Systems. 2002; pp 849–856.

(61) McGibbon, R. T.; Beauchamp, K. A.; Harrigan, M. P.; Klein, C.; Swails, J. M.; Hernández, C. X.; Schwantes, C. R.; Wang, L.-P.; Lane, T. J.; Pande, V. S. MD-Traj: A Modern Open Library for the Analysis of Molecular Dynamics Trajectories. Biophysical Journal 2015, 109, 1528 – 1532.

(62) Pedregosa, F.; Varoquaux, G.; Gramfort, A.; Michel, V.; Thirion, B.; Grisel, O.; Blondel, M.; Prettenhofer, P.; Weiss, R.; Dubourg, V., et al. Scikit-learn: Machine learning in Python, the Journal of machine Learning research 2011, 12, 2825–2830.

(63) Gulay, S. P.; Bista, S.; Varshney, A.; Kirmizialtin, S.; Sanbonmatsu, K. Y.; Dinman, J. D. Tracking fluctuation hotspots on the yeast ribosome through the elongation cycle. Nucleic Acids Research 2017, 45, 4958 4971.

(64) Kirmizialtin, S.; Hennelly, S. P.; Schug, A.; Onuchic, J. N.; Sanbonmatsu, K. Y. Integrating molecular dynamics simulations with chemical probing experiments using SHAPE-FIT. Methods in Enzymology 2015, 553, 215–234.

(65) Mardoum, W. M.; Gorczyca, S. M.; Regan, K. E.; Wu, T.-C.; Robertson-Anderson, R. M. Crowding Induces Entropically-Driven Changes to DNA Dynamics That Depend on Crowder Structure and Ionic Conditions. Frontiers in Physics 2018, 6, 53.

(66) Wilkinson, K. A.; Gorelick, R. J.; Vasa, S. M.; Guex, N.; Rein, A.; Mathews, D. H.; Giddings, M. C.; Weeks, K. M. High-Throughput SHAPE Analysis Reveals Structures in HIV-1 Genomic RNA Strongly Conserved across Distinct Biological States. PLoS Biology 2008, 6, e96.

(67) Aboul-ela, F.; Karn, J.; Varani, G. Structure of HIV-1 TAR RNA in the Absence of Ligands Reveals a Novel Conformation of the Trinucleotide Bulge. Nucleic Acids Research 1996, 24, 3974–3981.

(68) Kompella, V. P. S.; Romano, M. C.; Stansfield, I.; Mancera, R. L. Diffusion properties of transfer RNAs in the yeast cytoplasm under normal and osmotic stress conditions. Biochimica et Biophysica Acta (BBA) - General Subjects 2025, 1869, 130798.

(69) Chapman, C. D.; Gorczyca, S.; Robertson-Anderson, R. M. Crowding induces complex ergodic diffusion and dynamic elongation of large DNA molecules. Biophysical Journal 2015, 108, 1220–1228.

(70) Nawrocki, G.; Wang, P.-H.; Yu, I.; Sugita, Y.; Feig, M. Slow-Down in Diffusion in Crowded Protein Solutions Correlates with Transient Cluster Formation. The Journal of Physical Chemistry B 2017, 121, 11072 11084.

(71) Chen, Z.-J.; Tan, Z.-J. Predicting molecular crowding effects in ion-RNA interactions. Biophysical Journal 2018, 114, 2661–2673.

(72) McConkey, E. H. Molecular evolution, intracellular organization, and the quinary structure of proteins. Proceedings of the National Academy of Sciences 1982, 79, 3236–3240.

(73) Rouskin, S.; Zubradt, M.; Washietl, S.; Kellis, M.; Weissman, J. S. Genome-wide probing of RNA structure reveals active unfolding of mRNA structures in vivo. Nature 2014, 505, 701 705.

(74) Ben-Yaakov, D.; Andelman, D.; Podgornik, R. Dielectric decrement as a source of ion-specific effects. The Journal of Chemical Physics 2011, 134, 074705.

(75) Rohs, R.; West, S. M.; Sosinsky, A.; Liu, P.; Mann, R. S.; Honig, B. The role of DNA shape in protein–DNA recognition. Nature 2009, 461, 1248 1253.

(76) Nolan, V.; Sánchez, J. M.; Perillo, M. A. PEG-induced molecular crowding leads to a relaxed conformation, higher thermal stability and lower catalytic efficiency of Escherichia coli β-galactosidase. Colloids and Surfaces B: Biointerfaces 2015, 136, 1202–1206.

(77) de Vrieze, J. Pfizer’s vaccine raises allergy concerns. Science 2021, 371, 10–11.

(78) Moon, J. D.; Webber, T. R.; Brown, D. R.; Richardson, P. M.; Casey, T. M.; Segalman, R. A.; Shell, M. S.; Han, S. Nanoscale water–polymer interactions tune macroscopic diffusivity of water in aqueous poly(ethylene oxide) solutions. Chemical Science 2024, 15, 2495–2508.

