## Supporting Information for "Protein Crowders Remodel RNA Electrostatics, Hydration, and Dynamics: A Challenge to Steric Crowding Models"

### Electrostatic Remodeling, Dehydration, and Dynamics Slowing of RNA by Protein Crowders: Beyond Volume Exclusion

### Supplementary Movies

**Movie S1:** Cross section of the 1-$\mu$s long simulation trajectory of an RNA molecule in a non-crowded environment. Frames are taken every 8 ps.

**Movie S2:** Cross section of the first 1.216 $\mu$s of the simulation trajectory of an RNA molecule in a PEG-crowded environment. Frames are taken every 8 ps.

**Movie S3:** Cross section of the first 1.216 $\mu$s of the simulation trajectory of an RNA molecule in a protein-crowded environment. Frames are taken every 8 ps.

**Movie S4:** RNA conformation in different environments. RNA in non-crowded conditions (N-RNA), in the presence of PEG crowders (G-RNA), or protein crowders (P-RNA) is shown in gray, orange, and blue, respectively. RNA structures were extracted every 4 ps from the non-crowded simulation trajectory and every 8 ps from the crowded simulation trajectories. A total of 251 RNA structures from the three different environments were combined into separate frames.

### Supplementary Tables

Table 1: Proportion of each protein residue in the protein-crowded simulation box.

| Residue | Total number of residues | Total number of atoms |
|---|---|---|
| ARG | 286 | 6864 |
| HIS | 102 | 1734 |
| LYS | 300 | 6608 |
| ASP | 228 | 2736 |
| GLU | 274 | 4110 |
| SER | 160 | 1760 |
| THR | 208 | 2912 |
| ASN | 155 | 2170 |
| GLN | 119 | 2023 |
| CYS | 52 | 577 |
| GLY | 333 | 2331 |
| PRO | 127 | 1778 |
| ALA | 257 | 2573 |
| VAL | 277 | 4432 |
| ILE | 279 | 5301 |
| LEU | 282 | 5358 |
| MET | 108 | 1874 |
| PHE | 141 | 2820 |
| TYR | 130 | 2733 |
| TRP | 31 | 744 |
| In total | 3849 | 61438 |

### Supplementary Figures

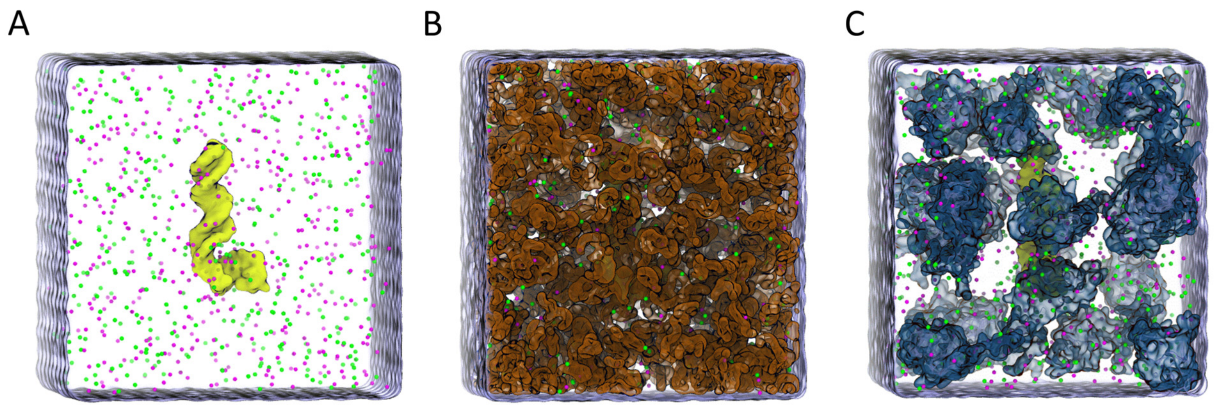


**Figure S1:** Overview of simulation-box configurations employed in this study to investigate RNA molecules within three distinct environments: A) non-crowded, B) PEG-crowded, and C) protein-crowded. Each simulation box contains the same RNA hairpin structure at its center. RNA, $Na^+$ ions, $Cl^-$ ions, PEG, and proteins are highlighted in yellow, purple, green, orange, and blue, respectively.

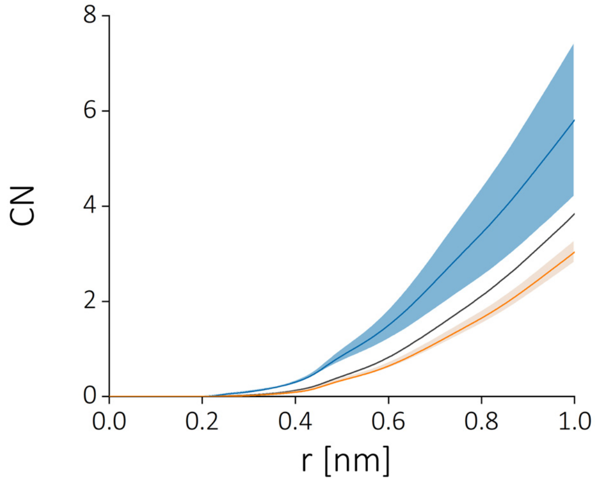


**Figure S2:** Impact of the environment on Cl- ion coordination. The coordination numbers (CN) of Cl- ions are provided for various distances from the RNA surface in non-crowded (dark grey), PEG-crowded (orange), and protein-crowded environments (blue).

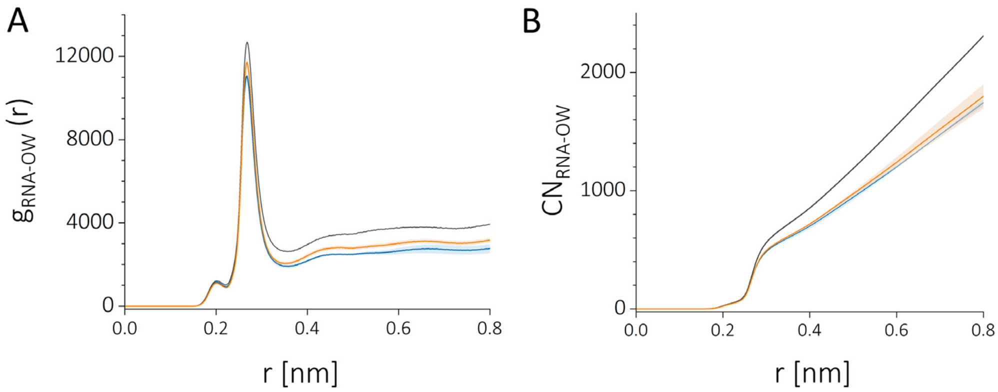


**Figure S3:** Crowding effects on RNA-water interactions. A, Surface RDF and, B, coordination number (CN) of water oxygen with RNA in non-crowded (dark grey), PEG-crowded (orange), and protein-crowded (blue) environments. Data derived from one non-crowded and two crowded simulation trajectories per condition.

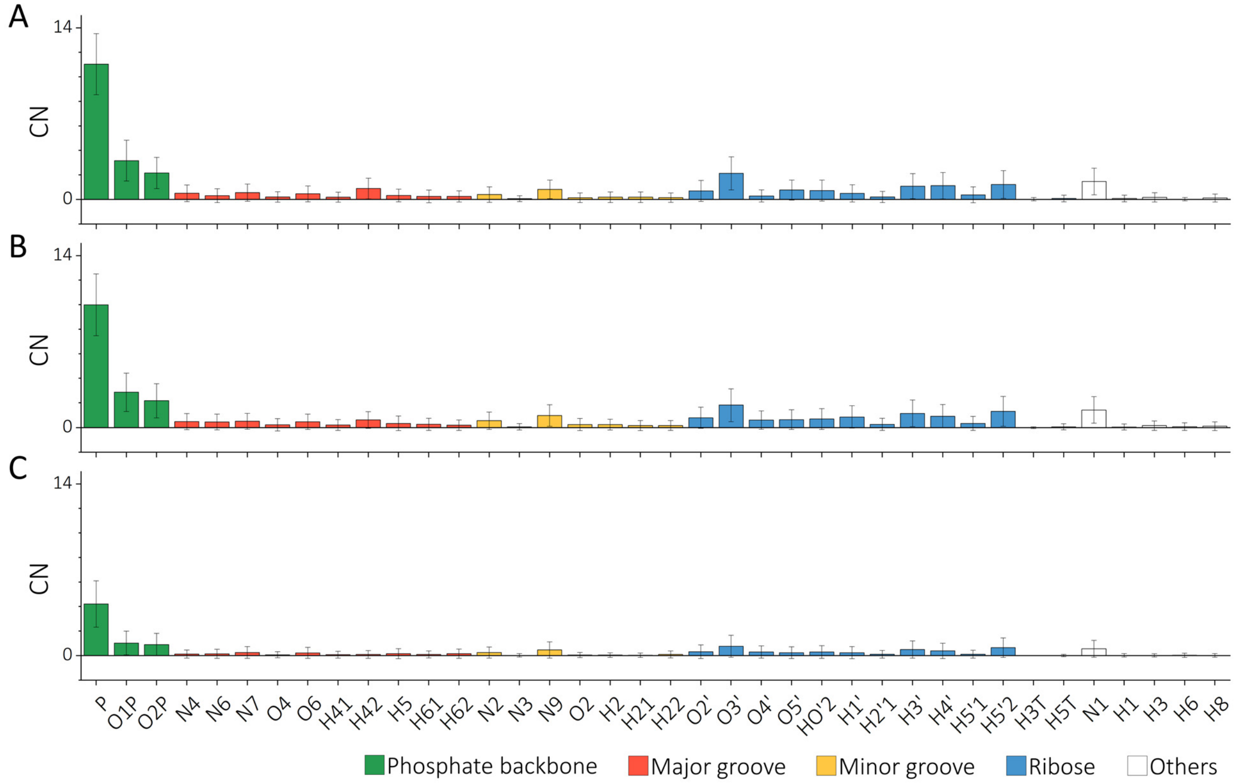


**Figure S4:** $Na^+$ ion binding pattern within 6 Å from the RNA surface in, A, a non-crowded, B, PEG-crowded, and, C, protein-crowded environment.

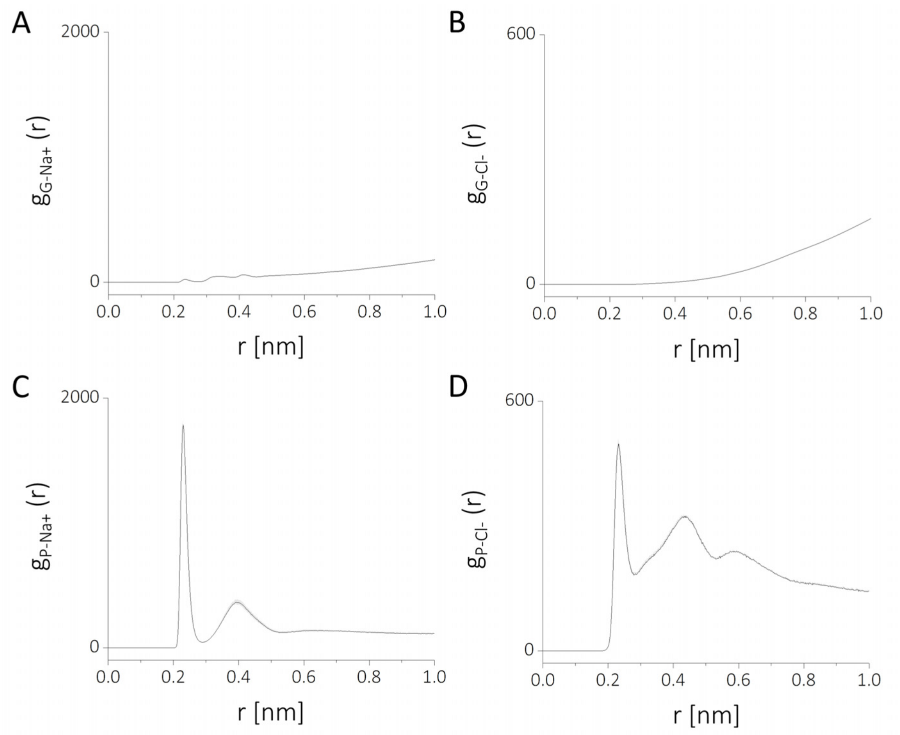


**Figure S5:** Comparison of interactions between crowding agents and ions. Surface RDF of, A, $Na^+$ ions with PEG, B, $Cl^-$ ions with PEG, C, $Na^+$ ions with proteins, and, D, $Cl^-$ ions with proteins.

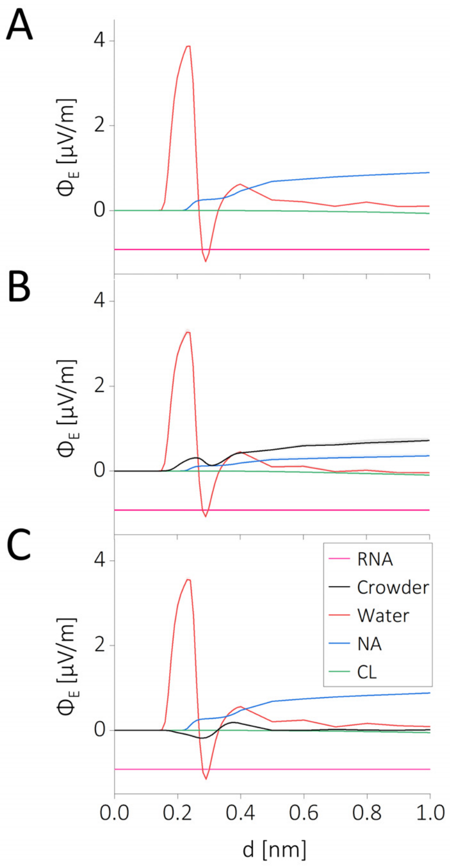


**Figure S6:** The influence of the environment on electric flux within a distance, d, from the RNA surface. Comparison of electrostatic flux resulting from various solute and solvent components among the non-crowded (A), PEG-crowded (B), and protein-crowded environments (C).

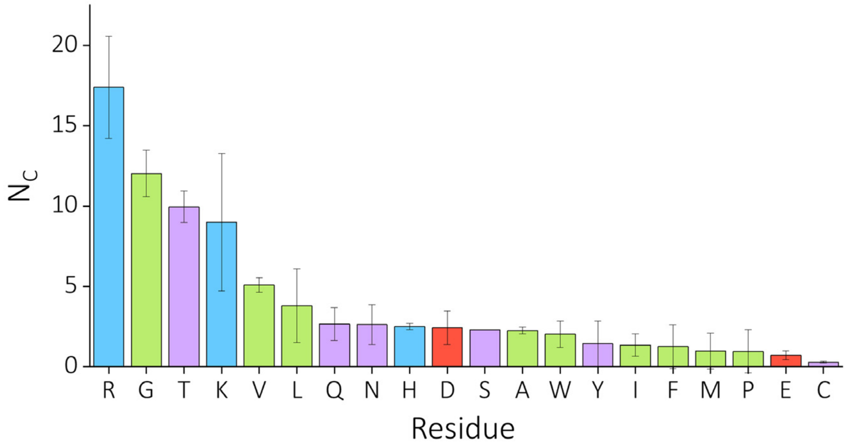


**Figure S7:** Amino acid binding frequency to the RNA events for a 6 Å cutoff.

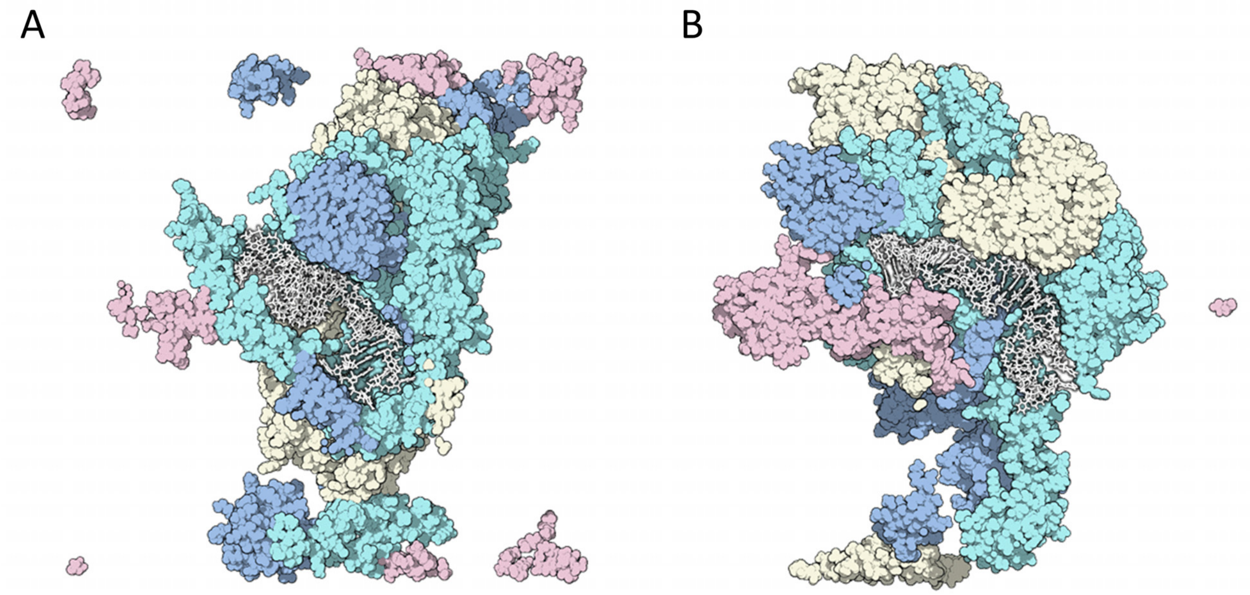


**Figure S8:** Comparison of protein distribution around the RNA in high-crowded (A) and low-crowded (B) environments. The figures depict cross-sectional views of the simulation box, with RNA represented in white using a 'licorice' style and proteins displayed as surfaces. Proteins S18, TCTP, L23, and Tα are highlighted in light blue, blue, light pink, and light yellow, respectively.

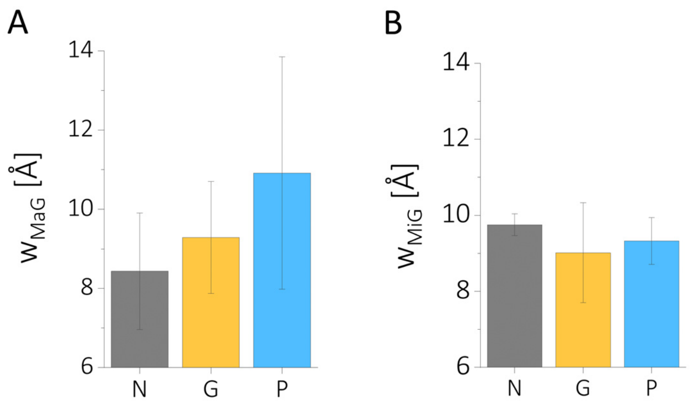


**Figure S9:** Impact of the environment on the width of, A, the major groove ($w_{MAG}$) and, B, the minor groove ($w_{MiG}$) of RNA. Results for non-crowded, PEG-crowded, and protein-crowded environments are distinguished by dark grey, orange, and blue shades, respectively.
